# Partial coupling of the proliferation and differentiation programs during *C. elegans* intestine development

**DOI:** 10.1101/2024.04.07.588410

**Authors:** Joris Dieng, Harpreet Singh, Grégoire Michaux, Anne Pacquelet

## Abstract

Cell proliferation and differentiation are essential processes underlying multicellular organism development. Cell proliferation arrest usually precedes terminal differentiation, suggesting that these two processes may be coordinated. Here we took advantage of the very stereotyped development of the *C. elegans* intestine to address whether the control of the proliferation and differentiation programs are systematically coupled. We show that delaying cell cycle arrest does not affect most aspects of intestinal differentiation but leads to a specific delay in the accumulation of late microvilli components. Reciprocally, we find that the differentiation factors ELT-2 and ELT-7 control cell cycle arrest specifically in posterior enterocytes. The occurrence of supernumerary divisions in the absence of ELT-2 and ELT-7 is associated with changes in the expression pattern of the cell cycle regulators cyclin B1 and CKI-1 and depends on the presence of the posterior Hox protein PHP-3. Our work thus demonstrates the existence of reciprocal interactions between cell proliferation and cell differentiation. It nevertheless also shows that these two processes are only partially coupled, suggesting the existence of additional mechanisms ensuring their temporal control.

## Introduction

The development of a single-cell egg into a multicellular organism involves both the generation of a large number of cells through cell proliferation and the acquisition of specialized cell functions through differentiation. Cell proliferation arrest usually precedes terminal differentiation, suggesting the existence of mechanisms that coordinate the two processes. Consistently, several mechanisms through which cell cycle regulators control the activity of differentiation factors have been described. Reciprocally, differentiation factors have been shown to induce the expression of cell cycle inhibitors (Ruijtenberg and van den Heuvel, 2016). The inverse correlation between proliferation and differentiation has also led to the common hypothesis that cell cycle exit may be a prerequisite to differentiation. Although this is indeed the case in some instances (e.g. (Willms et al., 2020)), the situation appears to be more complex than anticipated and some examples of uncoupling between cell cycle exit and differentiation have been described (Cockburn et al., 2022; Lobjois et al., 2008; Martinez et al., 2024). We therefore decided to investigate further how changes in cell proliferation affect differentiation during development and, reciprocally, how differentiation contributes to cell cycle regulation.

To address this question, we took advantage of the extremely stereotyped and well characterized development of the *C. elegans* intestine. This simple intestine is composed of 20 cells, which all derive from a single embryonic precursor cell, the E blastomere. This blastomere undergoes a series of cell divisions that successively give rise to the so-called E2, E4, E8, E16 and E20 intestinal development stages (Fig. S1A). First, the E blastomere and all its descendants divide four times to produce 16 enterocytes, which are organized into eight intestinal rings, each being formed by two cells. Cells from the first and last intestinal rings then divide once more, thereby giving rise to the 20-cell stage intestine, formed by nine intestinal rings, all containing two cells, except the first anterior ring which is made of four cells (Asan et al., 2016; Leung et al., 1999; Sulston et al., 1983).

While all early embryonic divisions are characterized by a quick succession of S and M phases, cell cycle length progressively increases in the E lineage from the E2 stage on until it becomes markedly long between the E16 and E20 stage (Leung et al., 1999; Sulston et al., 1983). The initial increase in intestinal cell cycle length is due to the introduction of a G2 phase at the E2 stage while a G1 phase is probably introduced only at the E16 stage (Edgar and McGhee, 1988; Wong et al., 2022). The CDC-25.1 phosphatase, which promotes G2/M transition through the activation of the cyclin dependent kinase CDK-1, has been shown to be required for intestinal divisions and embryos carrying *cdc-25.1* gain-of-function mutations present an acceleration of intestinal divisions (Bao et al., 2008; Clucas et al., 2002; Demouchy et al., 2024; Kostić and Roy, 2002). The CDC-25.2 phosphatase also appears to regulate intestinal divisions: in *cdc-25.2* loss-of-function mutants, the E2 to E4 division has been reported to be slower and the E16 to E20 division does not occur, suggesting that CDC-25.2 contributes to cell cycle progression together with CDC-25.1 in early intestinal divisions and becomes essential for the E16 to E20 division (Lee et al., 2016; Nair et al., 2013). On the other hand, the WEE-1.1 kinase starts to be expressed in the intestinal lineage at the E2 stage and prevents G2/M transition through the inhibition of CDK-1, thereby contributing to the increase of cell cycle length (Robertson et al., 2014). Consistent with the late appearance of a G1 phase, cyclin D1 (CYD-1), which promotes G1 progression, is required solely for the E16 to E20 division (Boxem and Van Den Heuvel, 2001). Loss of the CDK inhibitor CKI-1 also does not affect cell cycle length until the E16 stage but leads to the abnormal late division of all enterocytes after the E16 stage (Kostić and Roy, 2002). Altogether, those observations suggest that both CYD-1 and CKI-1 are essential regulators of the E16 to E20 division, with CYD-1 promoting G1 progression and division in the most anterior and posterior intestinal cells and CKI-1 preventing cell cycle progression in the remaining enterocytes.

Intestinal fate specification relies on the restricted expression of two GATA transcription factors, END- 1 and END-3, in the E blastomere (Maduro et al., 2005; Zhu et al., 1997; Zhu et al., 1998). END-1 and END-3 then trigger the expression of the differentiation GATA factors ELT-7 and ELT-2, which start to be expressed at the late E2 and E8 stage, respectively, and persist in enterocytes until adulthood (Demouchy et al., 2024; Sommermann et al., 2010; Wiesenfahrt et al., 2016). Both ELT-2 and ELT-7 control the transcription of a variety of intestinal specific genes, including digestion enzymes, nutrient transporters and microvilli components (Dineen et al., 2018; McGhee et al., 2009; Williams et al., 2023). *elt-7* null mutants lack a discernable phenotype (Sommermann et al., 2010); *elt-2* null mutants display short and irregular microvilli and die as L1 larvae but nevertheless display a continuous intestinal lumen as well as typical gut granules (Fukushige et al., 1998). The lack of phenotype in the absence of ELT-7 and the persistence of some aspects of intestinal differentiation in the absence of ELT-2 is due to the partial redundancy between ELT-2 and ELT-7 (Dineen et al., 2018; Sommermann et al., 2010). Consistently, in larvae lacking both ELT-2 and ELT-7, large portions of the intestine show neither a lumen nor gut granules (Sommermann et al., 2010).

Polarization of the intestinal epithelium and formation of the brush border are two major steps of intestinal differentiation. Polarization becomes apparent at the E16 stage when the polarity proteins PAR-3, PAR-6 and PKC-3 start to accumulate at the apical membrane of enterocytes (Achilleos et al., 2010; Leung et al., 1999; Totong et al., 2007). Some microvilli components (hereafter referred to as “early” components), including the intestine specific isoform of actin ACT-5, the actin/membrane linker ERM-1 and the filamin FLN-2, also start to localize at the apical membrane during the E16 stage, prior to microvilli formation (Bidaud-Meynard et al., 2021). Microvilli start to grow between the 1.5- and 2-fold stage embryonic stages, when the intestine has reached the E20 stage (Asan et al., 2016; Bidaud-Meynard et al., 2021). At this point, the apical localization of ACT-5 and ERM-1 markedly increases and additional microvilli components (referred to as “late” components), in particular the actin-capping protein EPS-8 and the unconventional myosin HUM-5, are also recruited to growing microvilli (Bidaud-Meynard et al., 2021). Microvilli then continue to grow and form a typical regular brush border at the 3-fold stage (Bidaud-Meynard et al., 2021).

Here we examine the reciprocal interactions that may exist between intestinal differentiation and divisions. While most aspects of intestinal differentiation occur independently of cell cycle arrest, we show that delaying cell cycle arrest specifically delays the apical accumulation of late microvilli components. Reciprocally, we find that the lack of the differentiation factors ELT-2 and ELT-7 induce supernumerary intestinal divisions. Notably this phenomenon is restricted to the most posterior enterocytes and depends on the presence of the posterior Hox protein PHP-3.

## Results

### Acceleration of intestinal cell cycles does not lead to precocious polarization

In control embryos, enterocyte polarization occurs at the E16 stage, just before the beginning of embryonic elongation (Fig. S1A) (Achilleos et al., 2010; Leung et al., 1999; Totong et al., 2007). We first wondered whether the timing of this polarization process was defined by the number of cell cycles undergone by intestinal blastomeres. If this were the case, one would expect it to occur earlier in embryos in which intestinal divisions occur faster. To test this hypothesis, we took advantage of the *cdc-25.1(rr31)* gain-of-function mutation, which has been shown to accelerate intestinal cell cycles and to lead to an additional division of intestinal blastomeres (Demouchy et al., 2024; Kostić and Roy, 2002). As a consequence, *cdc-25.1(rr31)* embryos reach the E8 and E16 stage earlier than control embryos and eventually display an average of 38 enterocytes (Fig. S1A) (Demouchy et al., 2024; Kostić and Roy, 2002). Consistently, we observed that control embryos reached the E16 stage prior to the beginning of embryonic elongation whereas *cdc-25.1(rr31)* embryos already had 32 enterocytes at this same stage (Fig. S1B). To assess the timing of epithelial polarization, we examined the apical accumulation of the polarity protein PAR-3. As previously described, in control embryos, PAR-3 was visible on a few non-polarized puncta at the E8 stage, started to accumulate in discrete foci at the apical membrane of enterocytes at the early E16 stage and eventually formed a continuous apical line later during the E16 stage, when the embryo was about to start elongating (Fig. S1B) (Achilleos et al., 2010; Naturale et al., 2023). In *cdc-25.1(rr31)* embryos, a few non-polarized PAR-3 puncta were visible at the E16 stage while foci started to accumulate apically at the early E32 stage and formed a continuous line later during the E32 stage, short prior to the beginning of elongation (Fig. S1B). Thus, although intestinal cells divide faster in *cdc-25.1(rr31)* embryos, they do not polarize prematurely, indicating that reaching the E16 stage is not sufficient to trigger polarization.

### Delaying cell cycle arrest delays the apical accumulation of late microvilli components

We next asked whether cell cycle arrest was a prerequisite for the different aspects of intestinal differentiation. The CDK inhibitor CKI-1 was previously shown to be required for cell cycle arrest of central enterocytes once the intestine has reached the E16 stage (Fig. S2A) (Kostić and Roy, 2002). By using an auxin-inducible degron which specifically degrades CKI-1 in the intestine (see material and methods for details), we could induce the appearance of late supplemental intestinal divisions and test the effect of these abnormal divisions on intestinal differentiation. First, we performed time-lapse recordings to carefully characterize the timing of the last intestinal divisions in control and CKI- 1::degron embryos. In control embryos, the E16 stage was reached prior to embryonic elongation and divisions of the first and last intestinal ring cells, which give rise to the E20 stage, occurred between the lima bean and 1.5-fold stage (Fig. S2B). Consistent with previous studies, we found that in CKI- 1::degron embryos all enterocytes divided after the E16 stage; these supernumerary divisions also occurred between the lima bean and 1.5-fold stage (Fig. S2B). Quantification of the number of enterocytes in embryos at different elongation stages confirmed that the E16 to E20 divisions of control embryos and the supernumerary divisions of CKI-1::degron embryos occurred between the lima bean and 1.5-stage (Fig. S2C).

We next asked whether the delay in cell cycle arrest induced by CKI-1 degradation delays intestinal differentiation. We first examined the effect of CKI-1 degradation on the expression of the differentiation factor ELT-2. In control embryos, ELT-2::mNG progressively accumulated after the E16 stage, while the embryo elongated (Fig1A). This accumulation was similar in CKI-1::degron embryos (Fig1A). We then looked at the intestine specific lysosome-related organelles called “gut granules” (Hermann et al., 2005) and could first detect them at the E16 stage both in control and CKI-1::degron embryos (Fig.1B). Finally, we found that the apical accumulation of the polarity protein PAR-6 at the very beginning of elongation was also similar in control and CKI-1::degron embryos (Fig.1C). Thus, delaying cell cycle arrest does not delay ELT-2 accumulation, gut granule formation or polarization, indicating that intestinal cells are able to go through these essential differentiation steps even when they still divide.

**Fig. 1:**
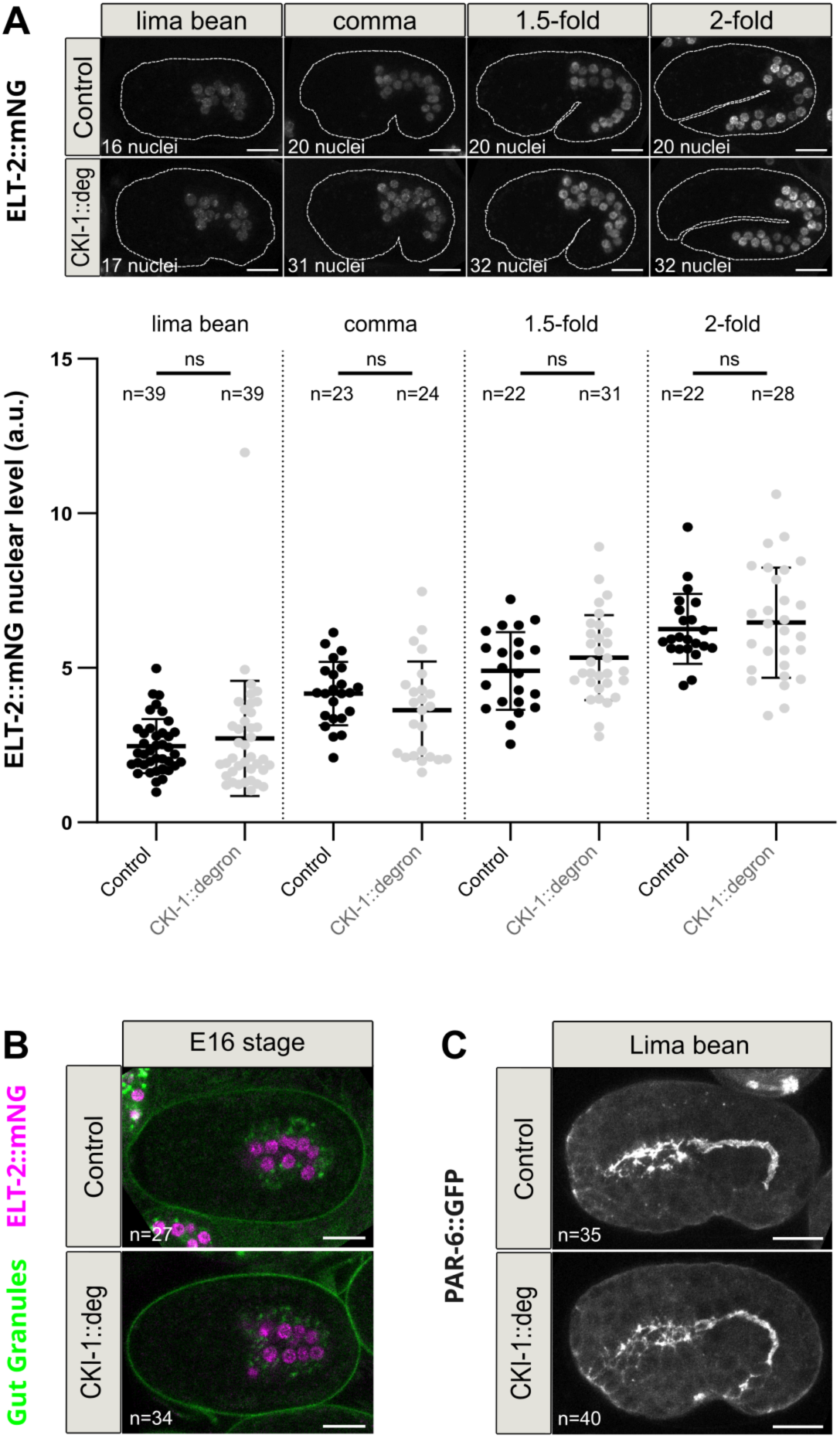
Delay of the cell cycle arrest induced by CKI-1 degradation does not delay most aspects of intestinal differentiation. **A.** Maximum Z-projections from confocal images of auxin-treated embryos expressing ELT-2::mNG and either *elt-2p::*TIR-1 (Control) or both *elt-2p::*TIR-1 and CKI-1::degron::BFP (CKI-1::deg) at the indicated embryonic stages and quantification of ELT-2::mNG nuclear fluorescence levels in those embryos (a.u.: arbitrary unit). ns: non-significant (*P*>0.05). Mann-Whitney test for lima bean and 2-fold stages, Welch’s t test for comma, unpaired t test for 1.5-fold. **B.** Maximum Z-projections from confocal images of auxin-treated E16 embryos expressing ELT-2::mNG (magenta) and either *elt-2p::*TIR-1 (Control) or both *elt-2p::*TIR-1 and CKI-1::degron::BFP (CKI-1::deg). Autofluorescent gut granules are visible in green. **C.** Maximum Z-projections from confocal images of auxin-treated lima bean embryos expressing PAR-6::GFP and either *elt-2p::*TIR-1 (Control) or both *elt-2p::*TIR-1 and CKI-1::degron::BFP (CKI-1::deg). Scale bars: 10 µm.

We then analyzed the effect of CKI-1 degradation on the apical accumulation of microvilli components. We first looked at two proteins known to accumulate early during intestinal development, ERM-1 and the filamin FLN-2 (Bidaud-Meynard et al., 2021). In control embryos, ERM-1 and FLN-2 already accumulated apically at the beginning of elongation (Fig2A-B). This apical accumulation was not delayed upon CKI-1::degradation (Fig2A-B). There was even a slight increase in FLN-2 apical level at the 1.5 and 2-fold stages (Fig.2B). We then examined the apical accumulation of two “late” microvilli components, namely EPS-8 and HUM-5 (Bidaud-Meynard et al., 2021). In control embryos, apical EPS-8 and HUM-5 started to be weakly detected at the 1.5-fold stage, became clearly apparent at the 2-fold stage and markedly accumulated at the 3 and 4-fold stages (Fig.2C-D). Notably, the pattern of EPS-8 localization also changed between the 1.5 and 2-fold stages: while it displayed a junction-like pattern at the 1.5-fold stage, it formed a continuous thicker apical line at the 2-fold stage (Fig.2C, insets). By contrast, following CKI-1 degradation, EPS-8 and HUM-5 apical levels remained very low at the 2-fold stage and started to increase only at the 3-fold stage (Fig.2C-D). EPS-8 also retained its junction-like pattern until the 2-fold stage (Fig.2C, inset). These observations strongly suggest that the late apical accumulation of EPS-8 and HUM-5 requires prior arrest of the cell cycle, thereby pointing towards the existence of a mechanism linking the timing of this aspect of intestinal differentiation to cell cycle progression.

**Fig. 2:**
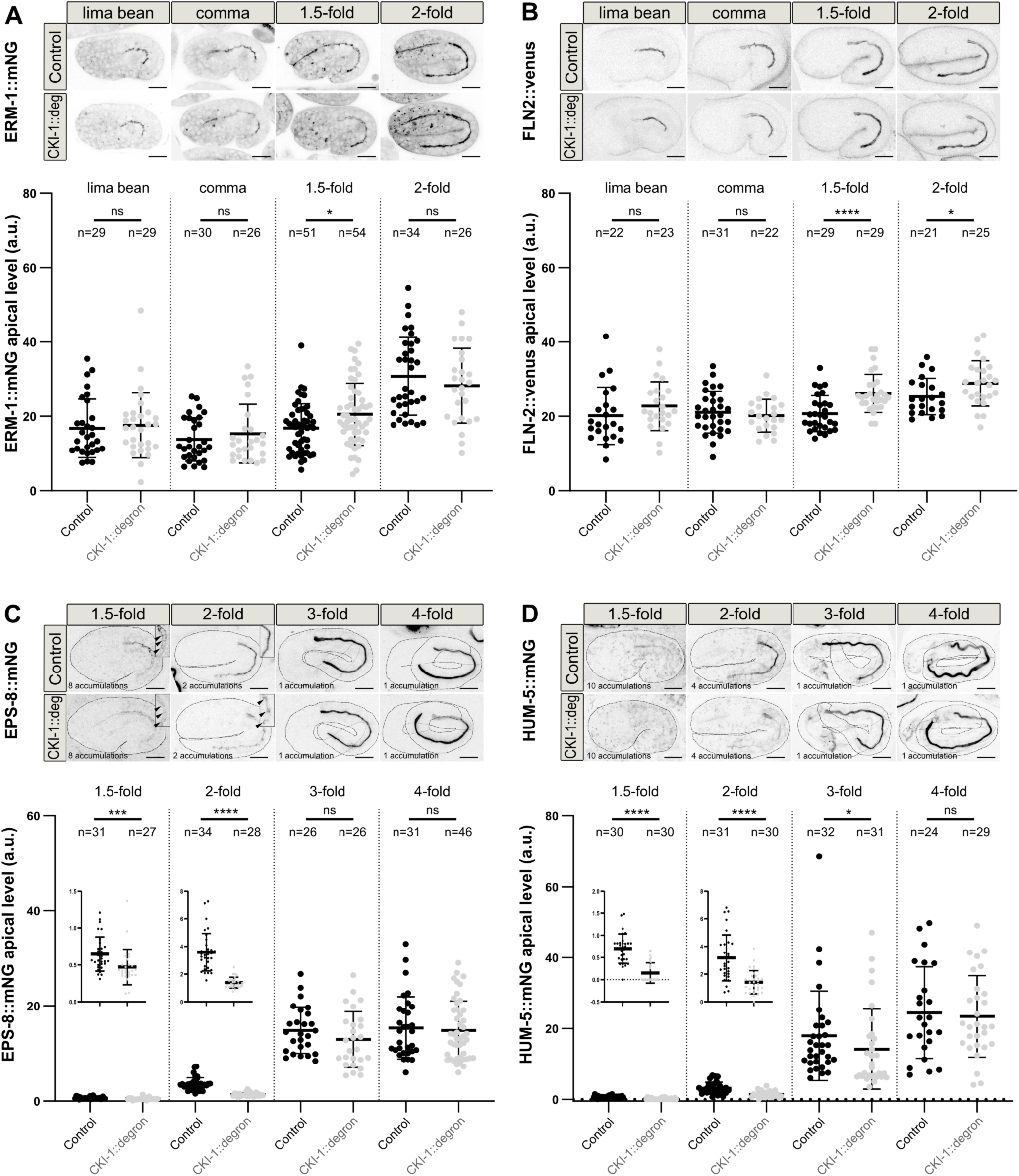
Delay of the cell cycle arrest induced by CKI-1 degradation delays the apical accumulation of late microvilli components. Maximum Z-projections from confocal images of auxin-treated embryos expressing ERM-1::mNG (A), FLN-2::venus (B), EPS-8::mNG (C) or HUM-5::mNG (D) and either *elt-2p::*TIR-1 (Control) or both *elt-2p::*TIR-1 and CKI-1::degron::BFP (CKI-1::deg) at the indicated embryonic stages and quantification of ERM-1::mNG (A), FLN-2::venus (B), EPS-8::mNG (C) and HUM-5::mNG (D) apical fluorescence levels in those embryos (a.u.: arbitrary unit). In C, insets on the top right corner of 1.5 and 2-fold embryos show a higher magnification of the intestinal apical region and highlight the EPS-8 junction-like pattern (arrowheads) observed in 1.5-fold embryos (control and CKI-1::degron) and in the 2-fold CKI-1::degron embryo. In C and D, the number of accumulations used for imaging the different embryonic stages are indicated (see materials and methods for details). Black dashed lines highlight embryo shape; at the 3- and 4-fold stages, fainter lines indicate the embryonic region which is folding in a lower focal plane. Magnifications of the graph area corresponding to 1.5-fold and 2-fold stages are shown to help visualizing changes in apical signal. In A to D, ns: non-significant (*P*>0.05), \**P*<0.05, \*\**P*<0.01, \*\*\**P*<0.001, \*\*\*\**P*<0.0001. In A, Mann-Whitney test for all stages; In B, unpaired t test for lima bean, comma and 2-fold, Mann-Whitney test for 1.5-fold; In C, Mann-Whitney test for 1.5-fold, 2-fold and 4-fold, unpaired t test for 3-fold; In D, Mann-Whitney test for 1.5-fold and 3-fold, Welch’s t test for 2-fold, unpaired t test for 4-fold. Scale bars: 10 µm.

### Intestinal polarization and brush border formation are not required to arrest intestinal cell divisions

We next sought to determine whether intestinal differentiation reciprocally controls cell cycle progression. We first tested whether the occurrence of two major differentiation steps, polarization and brush border formation, could contribute to cell cycle arrest. We first tested the possible role of polarization by degrading the polarity protein PAR-3 in intestinal cells (Fig. S3A) and found that 2-fold embryos without PAR-3 had the same number of intestinal nuclei as control embryos (Fig. S3B). We next assessed the importance of brush border formation by depleting ACT-5 (Fig. S3C), the intestinal specific form of actin, which has been shown to be required for microvilli formation (MacQueen et al., 2005), and observed that 2-fold embryos lacking ACT-5 displayed 20 intestinal nuclei, similar to control embryos (Fig. S3D). Thus, cell cycle arrest occurs in enterocytes independently of the onset of polarization or brush border formation.

### ELT-2 and ELT-7 are required to prevent the late appearance of supernumerary divisions in posterior enterocytes

We next asked whether the differentiation factors ELT-2 and ELT-7 themselves could be involved in controlling cell cycle progression. To this end, we depleted ELT-2 by RNAi and used *elt-7(tm840)* null mutants. *elt-2(RNAi)* was sufficient to induce a strong depletion of ELT-2 (Fig. S4A) as well as brush border defects (Fig. S4B). While both control and *elt-7(tm840)* 2-fold embryos displayed 20 intestinal nuclei, *elt-2(RNAi)* and *elt-7(tm840); elt-2(RNAi)* 2-fold embryos displayed a modest and marked increase in the number of intestinal nuclei, respectively (Fig.3A). Importantly, we did not observe a significant increase in the number of intestinal nuclei in *elt-7(tm840); elt-2(RNAi)* embryos at the very beginning of elongation (Fig.3B, early lima bean stage), suggesting that the presence of extra nuclei at the 2-fold stage is due to late supernumerary divisions. To test whether the presence of supernumerary intestinal nuclei was indeed due to supernumerary cell divisions and not for instance to endomitosis, we next counted the number of enterocytes in embryos expressing both the intestinal nuclear marker *end1p*::H1::mCherry and the membrane marker *end1p*::GFP::CAAX. This showed that *elt-7(tm840); elt-2(RNAi)* 2-fold embryos did not display only supernumerary intestinal nuclei but also supernumerary intestinal cells (Fig.3C). Altogether our observations indicate that ELT-2 and ELT-7 act redundantly to prevent abnormal late divisions of enterocytes. Notably, the most posterior nuclei (Fig.3A) and the most posterior cells (Fig.3C) of *elt-7(tm840); elt-2(RNAi)* intestines appeared smaller than others, suggesting that extra divisions may occur in the posterior region of the intestine.

**Fig. 3:**
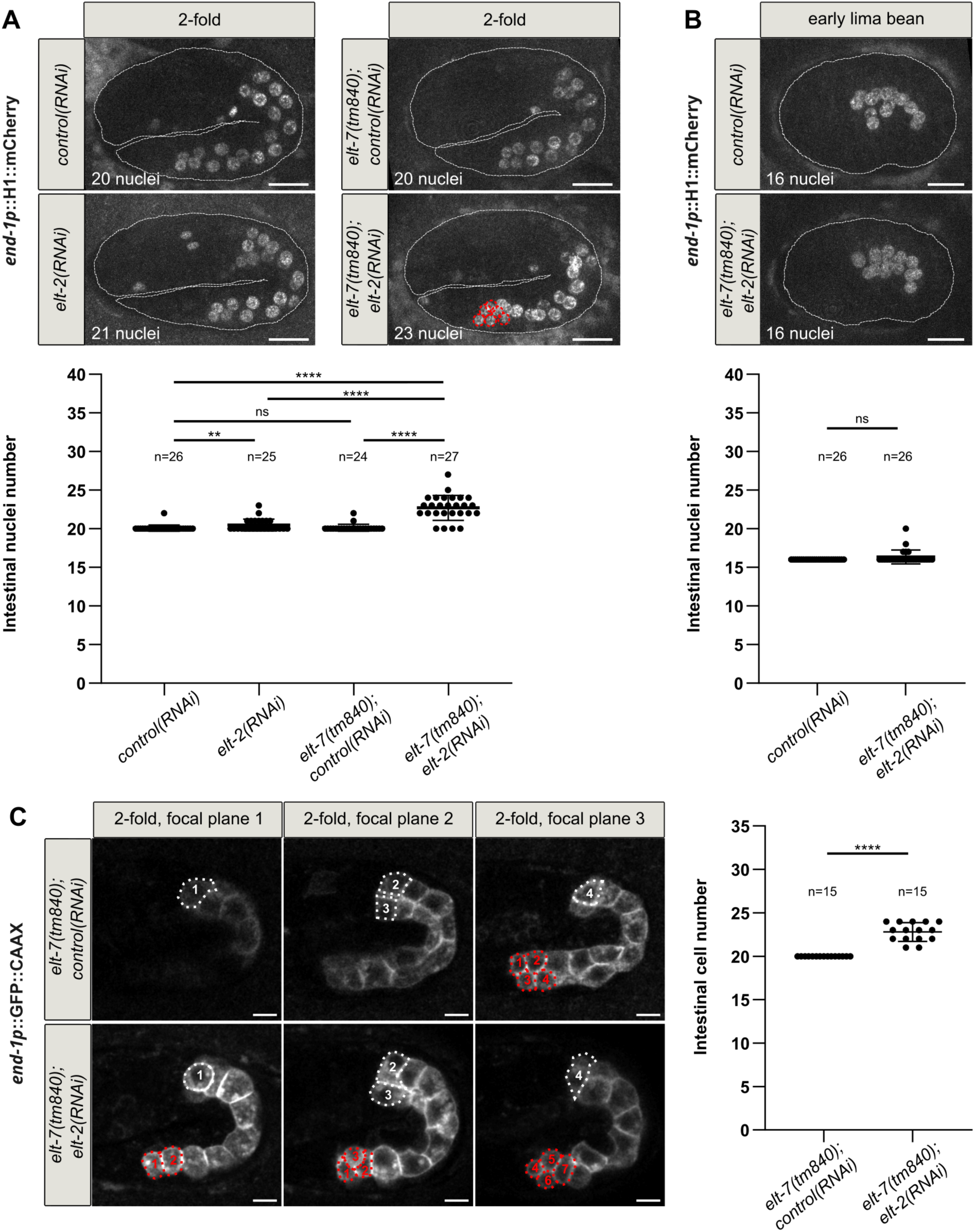
Loss of ELT-2 and ELT-7 leads to the presence of supernumerary enterocytes. **A.** Maximum Z-projections from confocal images of 2-fold control and *elt-7(tm840)* embryos expressing the intestinal nuclear marker *end1p::*H1::mCherry and treated with control or *elt-2* RNAi and quantification of the number of intestinal nuclei in those embryos. In *elt-7(tm840); elt-2(RNAi)* embryos, the posterior nuclei (red dotted circles) appear smaller than other intestinal nuclei, suggesting that they could result from supernumerary intestinal divisions. White dashed lines highlight embryo shape. ns: non-significant (*P*>0.05), \*\**P*<0.01, \*\*\*\**P*<0.0001, Mann-Whitney test. Scale bars: 10 µm. **B.** Maximum Z-projections from confocal images of early lima bean control and *elt-7(tm840)* embryos expressing *end1p::*H1::mCherry and treated with control or *elt-2* RNAi, respectively, and quantification of the number of intestinal nuclei in those embryos. White dashed lines highlight embryo shape. ns: non-significant (*P*>0.05), Mann-Whitney test. Scale bars: 10 µm. **C.** Confocal images taken at three different focal planes of *elt-7(tm840)* 2-fold embryos expressing the intestinal membrane marker *end1p*::GFP::CAAX and treated with control or *elt-2* RNAi and quantification of the number of intestinal cells in those embryos. Dotted lines highlight cells from the most anterior (white) and posterior (red) intestinal rings. *elt-7((tm840)* embryos have 20 enterocytes, with 4 anterior and 4 posterior cells arising from the divisions occurring between E16 and E20 stage. The *elt-7((tm840); elt-2(RNAi)* embryo shown has 23 enterocytes, with 7 small posterior cells that may result from late supernumerary divisions. \*\*\*\**P*<0.0001, Mann-Whitney test. Scale bars: 5 µm.

To undoubtedly determine the origin of supernumerary intestinal cells, we next recorded the late steps of intestinal development in *elt-7(tm840); elt-2(RNAi)* embryos expressing *end1p*::H1::mCherry and *end1p*::GFP::CAAX. These movies revealed that *elt-7(tm840); elt-2(RNAi)* embryos first underwent the E16 to E20 transition as described in wildtype embryos (Asan et al., 2016), with cells from both the most anterior and most posterior rings dividing once (Fig.4). However, *elt-7(tm840); elt-2(RNAi)* embryos then also displayed supernumerary divisions (n=8/10 embryos), which occurred on average 58 min (min/max=36/93 min, n=8) after the E20 stage and emanated from cells of the 9^th^ intestinal ring (n=8/8 embryos with supernumerary divisions) and from the 8^th^ intestinal ring (n=5/8 embryos with supernumerary divisions) (Fig.4). Thus, once the intestine has reached the E20 stage, the differentiation factors ELT-2 and ELT-7 appear to be required to arrest cell divisions specifically in the most posterior enterocytes.

**Fig. 4:**
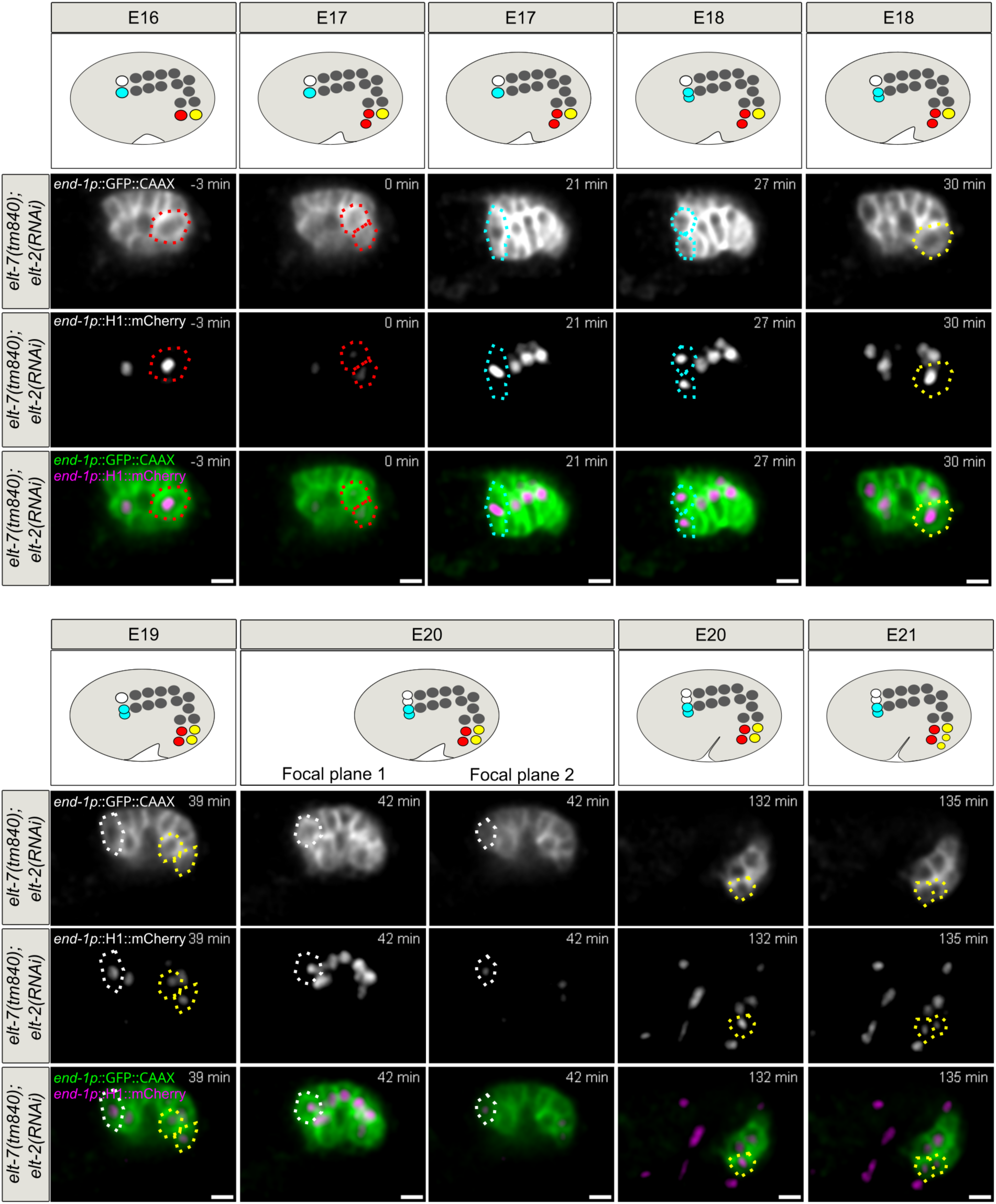
Loss of ELT-2 and ELT-7 leads to supernumerary late divisions of posterior enterocytes. Schematic representations and confocal images showing the enterocytes of a *elt-7(tm840)* embryo treated with *elt-2* RNAi, expressing the intestinal membrane marker *end1p*::GFP::CAAX and the intestinal nuclear marker *end1p::*H1::mCherry and recorded between the E16 and E21 stage. White/cyan and red/yellow circles highlight the enterocytes from the first and last intestinal ring, respectively, before (E16) and after (E17 to E21) their division. The first division of the E16 to E20 transition was used as reference for division timing (t=0). In this embryo, first ring divisions occur between t=21 to t=27 min (cyan cells) and t=39 to t=42 min (white cells; at t=42 min 2 focal planes are shown to make the two white daughter cells visible). Last ring divisions occur at t=0 (red cells) and between t=30 to t=39 min (yellow cells). Later, between t=132 to t=135, one of the yellow daughter cells undergoes a supernumerary division. Scale bars: 5 µm.

### Posterior enterocytes undergoing a supernumerary division show an abnormally late re-accumulation of Cyclin B1

We then set out to determine whether the appearance of a late additional division in the most posterior enterocytes was associated with changes in the expression of cell cycle regulators. First, we examined the expression of cyclin B1 (CYB-1), one of the three B-type cyclins, which, together with CDK-1, ensures proper progression through M phase (van der Voet et al., 2009). In E16 stage control embryos, CYB-1::GFP was never observed in cells that had stopped dividing but accumulated in the nuclei of enterocytes from the first and last intestinal rings before they divided to give rise to the E20 stage intestine (n=16/16 embryos) (Fig.S5). Occasionally, nuclear accumulation of CYB-1::GFP was also visible in one or two cells of the 7^th^ intestinal ring (n=5/16 and 2/16 embryos, respectively). These cells then also divided again, leading to the formation of an intestine composed of 21 or 22 cells. Such an occasional presence of 21 or 22 enterocytes has been previously described (Sulston and Horvitz, 1977). Importantly, CYB-1::GFP was no longer visible once the intestine had reached the E20 (or E21/E22) cell stage (Fig.S5). Similarly, in E16 stage *elt-7(tm840)* embryos treated with control RNAi, CYB-1::GFP accumulated in the nuclei of the first and last intestinal ring enterocytes before the E16 to E20 division and did not reappear in their daughter cells (n=12/12 embryos) (Fig.S6). In E16 stage *elt-7(tm840); elt-2(RNAi)* embryos, CYB-1::GFP also accumulated in the nuclei of the first and last intestinal ring enterocytes before the E16 to E20 division (n=11/11 embryos) (Fig.5). CYB-1::GFP then never reappeared in the daughter cells of the first intestinal ring (n=11/11 embryos) but accumulated again in one (n=4/11 embryos) or two (n=5/11 embryos) pairs of posterior daughter cells (Fig.5). Among these 9 embryos that displayed a late accumulation of CYB-1::GFP, 8 then underwent supernumerary late divisions. In 2/11 embryos, CYB-1::GFP did not reappear in the posterior daughter cells; no supernumerary division was observed in those embryos. Thus, in the absence of ELT-2 and ELT-7, the abnormally late expression of the mitotic cyclin CYB-1 precedes the supernumerary division of the most posterior enterocytes.

**Fig. 5:**
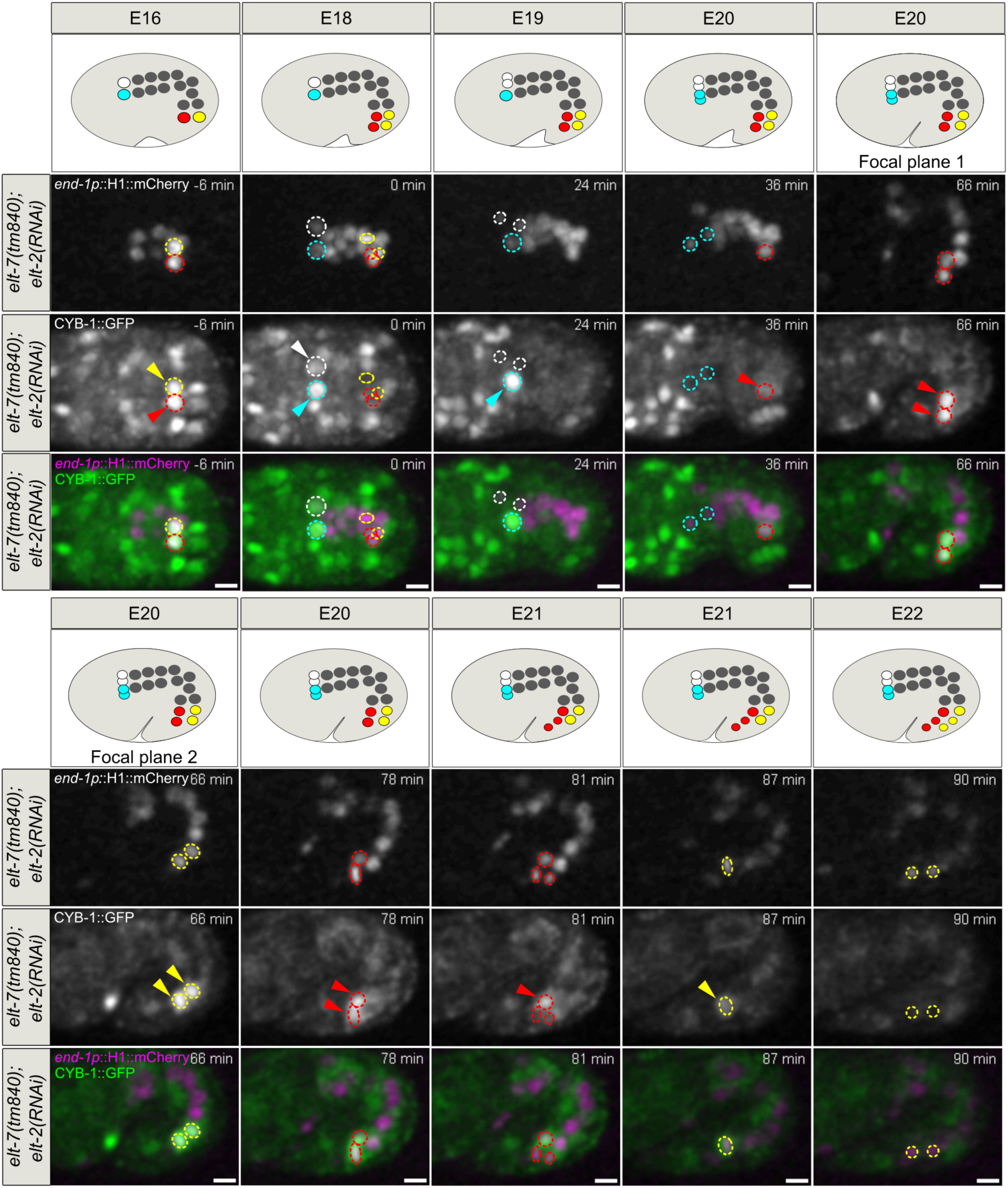
Loss of ELT-2 and ELT-7 is associated with the late re-accumulation of CYB-1::GFP in posterior enterocytes. Schematic representations and confocal images showing the enterocytes of a *elt-7(tm840)* embryo treated with *elt-2* RNAi, expressing the intestinal nuclear marker *end1p::*H1::mCherry and CYB-1::GFP and recorded between the E16 and E22 stage. White/cyan and red/yellow circles highlight the enterocytes from the first and last intestinal ring, respectively, before (E16) and after (E17 to E22) their division. The first division of the E16 to E20 transition was used as reference for division timing (t=0). In this embryo, first ring divisions occur at t=24 min (white nuclei) and t= 36 min (cyan nuclei). Last ring divisions occur at t=0 (red and yellow nuclei). CYB-1::GFP accumulates specifically in the first and last ring enterocytes prior to their division (t=-6 min, t=0 min and t=24 min, arrowheads) and does not reappear in the anterior daughter cells. At t=36 min, CYB-1::GFP starts to reappear in one cell of ring 9 (one of the red nuclei shown by arrowhead); at t=66 min, it is clearly visible in all cells from ring 8 and 9 (2 red and 2 yellow nuclei shown by arrowheads, 2 focal planes are shown to make the 4 nuclei visible). One of the red and one of the yellow daughter cells undergo a supernumerary division at t= 81 min and t=90 min, respectively. The other red and yellow daughter cells both undergo a supernumerary division immediately after (not shown). Scale bars: 5 µm.

### CKI-1 accumulation is delayed in posterior enterocytes undergoing a supernumerary division

We then examined whether the loss of ELT-2 and ELT-7 alters the expression of the cell cycle inhibitor CKI-1. During the E16/E20 transition, a transient accumulation of GFP::CKI-1 was observed in the most anterior and posterior enterocytes of control embryos (Fig. S7A-B). A weak nuclear signal sometimes preceded division: this signal was visible in 6/10 and 4/10 embryos for anterior and posterior cells, respectively, and appeared on average 20 min before division. GFP::CKI-1 then accumulated in the nuclei of anterior and posterior daughter cells just after division: 7/10 embryos displayed an accumulation in anterior nuclei (6 in both pairs of daughter cells, 1 in only one pair) and 9/10 in posterior nuclei (7 in both pairs of daughter cells and 2 in only one pair). GFP::CKI-1 levels then progressively decreased until no more GFP::CKI-1 could be detected (on average 35 and 40 minutes after the division of anterior and posterior cells, respectively). GFP::CKI-1 displayed a similar transient accumulation during the E16/E20 transition of *elt-7(tm840)* embryos treated with control RNAi (Fig. S8A-B): prior to the division of anterior and posterior cells, a weak nuclear signal was visible in n=8/10 and 4/10 embryos, respectively; after division, GFP::CKI-1 accumulated in the nuclei of anterior and posterior daughter cells (in all anterior daughter cell nuclei in 10/10 embryos, in all posterior daughter cell nuclei in 9/10 embryos and in one pair of posterior daughter cells in 1/10 embryo). In E16/E20 stage *elt-7(tm840); elt-2(RNAi)* embryos, GFP::CKI-1 also transiently accumulated in the two most anterior intestinal nuclei (Fig.6A): it was visible in 2/9 embryos before division and in 8/9 embryos after division (in all anterior daughter cells in 6/9 embryos and in one pair of anterior daughter cells in 2/9 embryos). By contrast, when the two most posterior enterocytes divided, GFP::CKI-1 was detected in 1/9 embryo prior to division and in none or only one pair of daughter cells after division in 5/9 and 4/9 embryos, respectively (Fig.6B). Among these 9 embryos, at least 8 then underwent late supernumerary divisions, with 7/8 showing a late nuclear accumulation of GFP::CKI-1 in supernumerary cells (Fig.6B). Our observations thus show that, during the E16/E20 transition, CKI-1 normally transiently accumulates in the nuclei of anterior and posterior enterocytes. This accumulation is likely to drive cell cycle arrest. However, in the absence of ELT-2 and ELT-7, the accumulation of CKI- 1 in posterior nuclei after the E16/E20 division is severely reduced; this lack of CKI-1 accumulation probably contributes to the ability of posterior cells to divide one more time.

**Fig. 6:**
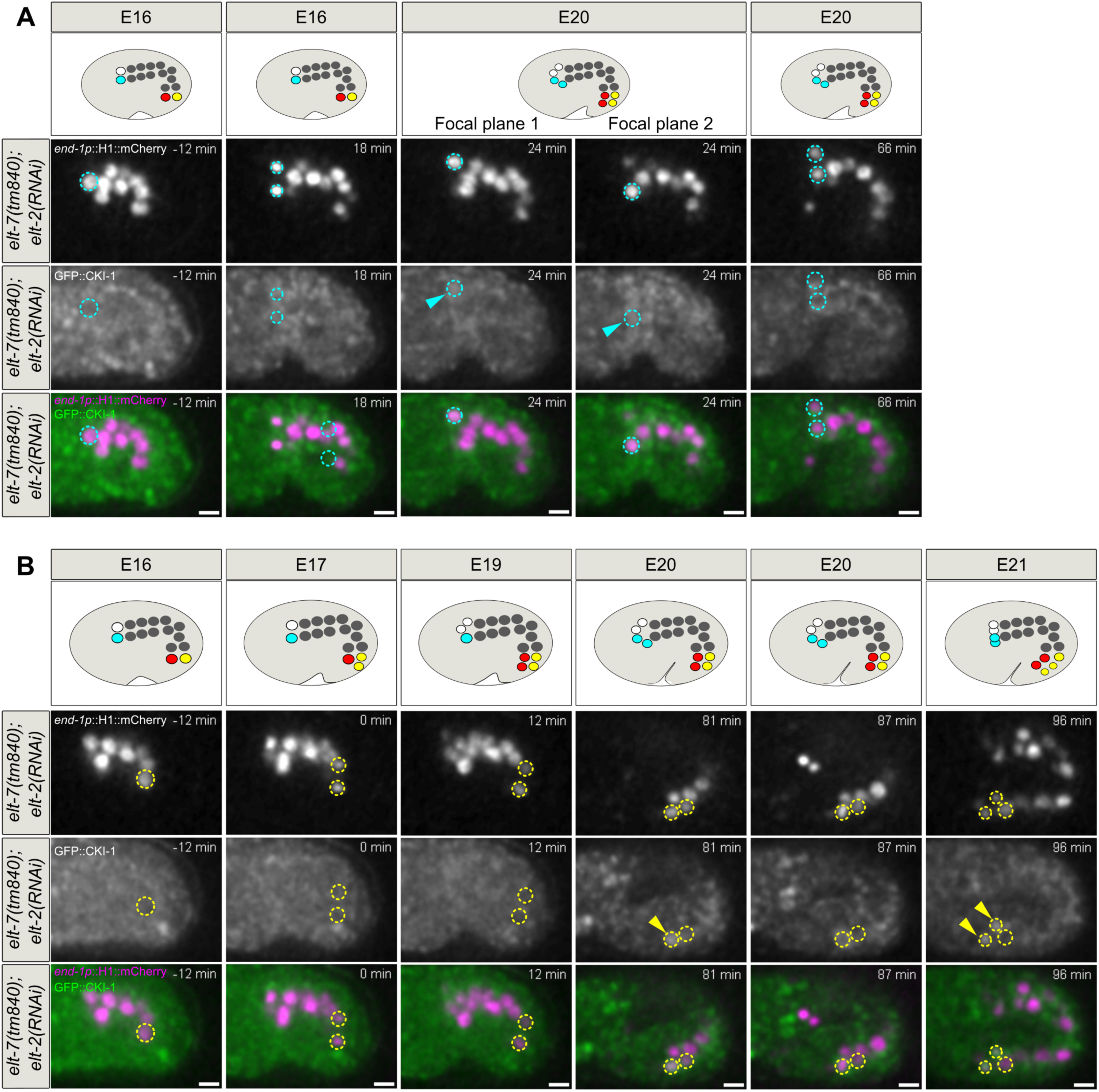
Loss of ELT-2 and ELT-7 is associated with loss of GFP::CKI-1 accumulation in posterior enterocytes. Schematic representations and confocal images showing the enterocytes of a *elt-7(tm840)* embryo treated with *elt-2* RNAi, expressing the intestinal nuclear marker *end1p::*H1::mCherry and GFP::CKI-1 and recorded between the E16 and E21 stage. The same embryo is shown in A and B; focal planes were chosen to display the anterior (A) or posterior (B) cell divisions. White/cyan and red/yellow circles highlight the nuclei of one cell of the first and last intestinal ring enterocytes, respectively, before (E16) and after (E17 to E21) their division. The first division of the E16 to E20 transition was used as reference for division timing (t=0). In this embryo, one first ring division (A) occurs at t=18 min (cyan nuclei). Division of the other anterior cell (white nucleus) occurs in a different focal plane and is not shown here. GFP::CKI-1 accumulates in the nucleus of the two cyan daughter cells after the division (t=24 min, arrowheads, 2 focal planes are shown to make the two daughter nuclei visible) and then progressively disappears until it is not any longer visible (t=66 min). One last ring division (B) occurs at t=0 (yellow nuclei). Division of the other posterior cell (red nucleus) occurs in a different focal plane and is not shown here. GFP::CKI-1 does not accumulate in the nucleus of the two yellow daughter cells right after the division (t=12 min). It eventually becomes visible in one of the two yellow daughters at t=81 min (arrowhead). This cell then undergoes a supernumerary division and its daughters then accumulate GFP::CKI-1 in the nucleus (t=96 min, arrowheads). Scale bars: 5 µm.

### Supernumerary divisions of posterior enterocytes depend on the presence of the posterior Hox protein PHP-3

We showed that ELT-2 and ELT-7 are required to prevent the occurrence of supernumerary late divisions specifically in posterior enterocytes. These observations suggest that posterior cells are more prone than other intestinal cells to escape cell cycle arrest once the intestine has reached the E20 stage. Posterior enterocytes distinguish themselves from more anterior cells in particular by the expression of two Hox transcription factors, NOB-1 and PHP-3 (Liu and Murray, 2023; Murray et al., 2022; Zhao et al., 2010). Consistently, we observed that both NOB-1::GFP and PHP-3::GFP were expressed in posterior enterocytes, respectively in rings 7 to 9 and 6 to 9, with PHP-3 being expressed at much higher levels than NOB-1::GFP (Fig.7A). We thus asked whether the expression of PHP-3 could explain the susceptibility of posterior enterocytes to undergo supernumerary divisions and investigated whether a *php-3* mutant could prevent the occurrence of these extra divisions. We found that *php-3(ok919); elt-7(tm840); elt-2(RNAi)* embryos showed a weaker increase in intestinal nuclei number compared to *elt-7(tm840); elt-2(RNAi)* embryos (Fig.7B), thus strongly suggesting that the presence of PHP-3 in posterior cell contributes to their ability to undergo supernumerary divisions in the absence of ELT-2 and ELT-7.

**Fig. 7:**
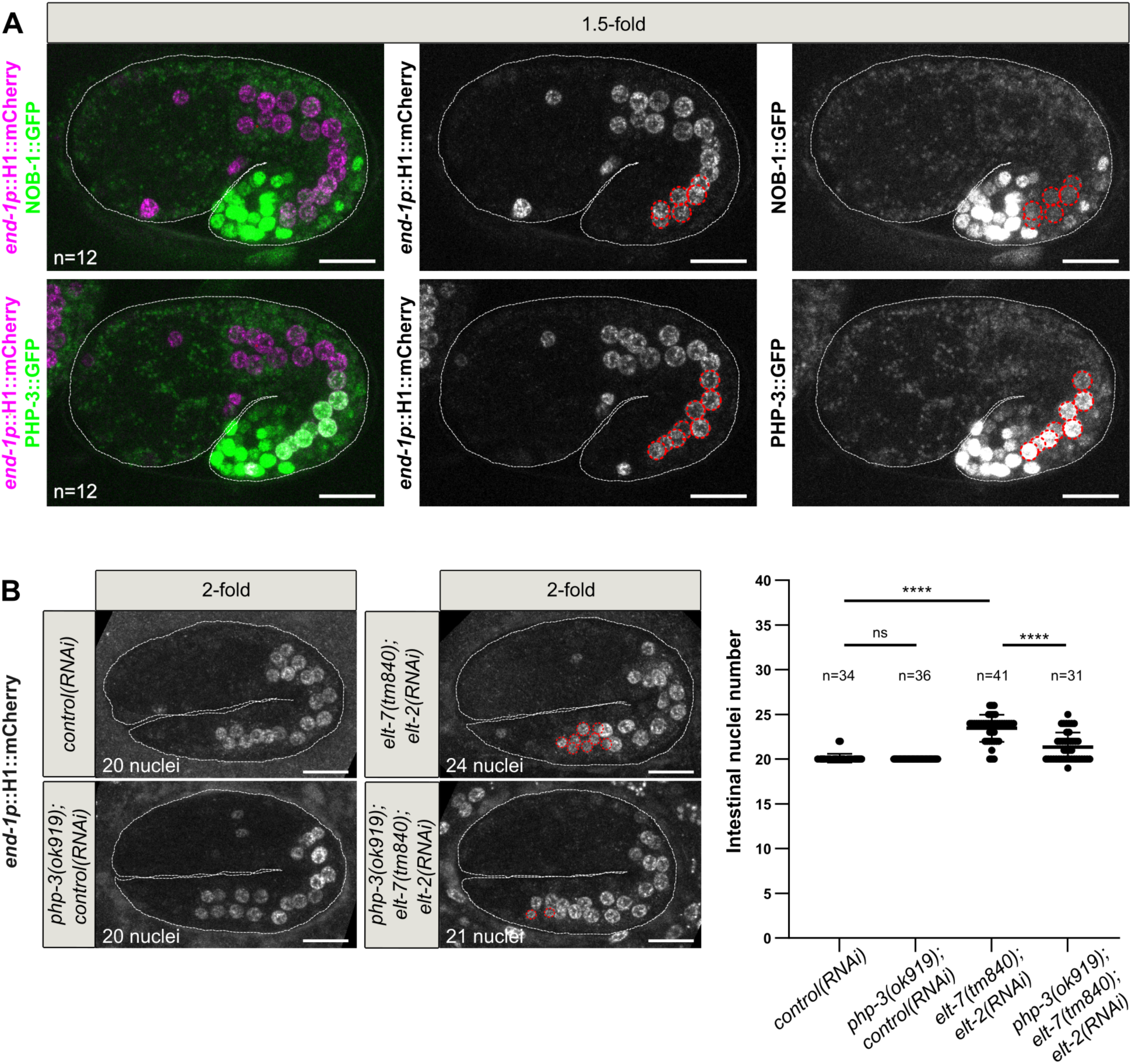
Supernumerary divisions of posterior enterocytes depend on the presence of the posterior Hox protein PHP-3. **A.** Maximum Z-projections from confocal images of 1.5-fold embryos expressing the intestinal nuclear marker *end1p::*H1::mCherry (magenta) and NOB-1::GFP (green, upper row) or PHP-3::GFP (green, lower row). NOB-1 and PHP-3 are expressed in posterior enterocytes (rings 7 to 9 for NOB-1::GFP, rings 6 to 9 for PHP-3::GFP, red dotted circles). **B.** Maximum Z-projections from confocal images of 2-fold control, *php-3(ok919)*, *elt-7(tm840)* and *php-3(ok919); elt-7(tm840)* embryos expressing *end1p::*H1::mCherry and treated with control or *elt-2* RNAi and quantification of the number of intestinal nuclei in those embryos. ns: non-significant (*P*>0.05), \*\*\*\**P*<0.0001, Mann-Whitney test. Red dotted circles highlight smaller posterior nuclei, which are likely to result from supernumerary intestinal divisions. In A and B, white dashed lines highlight embryo shape. Scale bars: 10 µm.

Altogether, our results demonstrate that ELT-2 and ELT-7 are required to prevent to occurrence of supernumerary late divisions in posterior intestinal cells. These supernumerary divisions are accompanied by changes in the accumulation of cell cycle regulators and depend on the presence of the posterior Hox protein PHP-3.

## Discussion

In this study, we addressed how cell proliferation and differentiation are regulated during development by using the *C. elegans* intestine as a model system. We have shown that acceleration of intestinal divisions does not impact the timing of polarization whereas delaying cell cycle arrest specifically delays the accumulation of late microvilli components. Reciprocally, we have also found that the differentiation factors ELT-2 and ELT-7 are required to arrest intestinal divisions specifically in the most posterior enterocytes.

The first signs of intestinal differentiation, including polarization, gut granule formation and apical accumulation of some microvilli components such as ERM-1 and FLN-2, are visible at the E16 stage, when most enterocytes have ceased dividing, during the long interphase that precedes the last division of the most anterior and posterior enterocytes (Achilleos et al., 2010; Bidaud-Meynard et al., 2021; Leung et al., 1999; Totong et al., 2007). The next important steps of differentiation, in particular microvilli growth and nutrient transporter expression, are initiated at the 2-fold and 3-fold stage, respectively, once the intestine has reached the E20 stage and all intestinal cells have undergone cell cycle arrest (Asan et al., 2016; Bidaud-Meynard et al., 2019). This led us to wonder whether the timing of intestinal cell divisions and differentiation were regulated by two independent timers that happen to coincide or whether some mechanisms exist to actively coordinate the two processes. In the latter case, we hypothesized that either the number of cell divisions undergone by enterocytes could be used as a clock controlling differentiation or, alternatively, that the cell cycle status, namely cell cycle arrest, could trigger differentiation. However, we find that acceleration of the cell cycle induced by a *cdc-25.1* gain-of-function mutation does not lead to premature enterocyte polarization. Thus, the number of divisions undergone by intestinal cells is not sufficient to provide the temporal information required to induce polarization in a timely manner. Interestingly, it was previously shown that slowing down early intestinal cell cycles does not delay the onset of *elt-2* transcription (Nair et al., 2013), further suggesting that the number of cell cycles undergone by intestinal cells is not a critical signal for their ability to start differentiating. Moreover, we also show that cell cycle arrest is not a prerequisite for early intestinal differentiation steps, namely for polarization, gut granule formation and accumulation of the early microvilli components ERM-1 and FLN-2. This suggests the existence of alternative mechanisms that control the timing of these differentiation processes. One possibility is that the progressive accumulation of the differentiation factor ELT-2, which is also independent of cell cycle arrest, provides an internal clock for enterocytes. By contrast, we show that degrading the cell cycle inhibitor CKI-1 is sufficient to delay the apical accumulation of the microvilli components EPS-8 and HUM-5. These observations indicate that CKI-1 is required to induce the timely accumulation of late microvilli components, thereby revealing its ability to couple cell cycle arrest with some aspects of intestinal differentiation. Altogether, our observations confirm that the relationships between cell cycle and differentiation are dependent on the developmental context. The ability of cells to initiate differentiation before cell cycle arrest may favor a smooth transition from undifferentiated progenitors to fully differentiated post-mitotic cells. It has also been proposed to help producing a sufficient pool of differentiated cells in tissues with a high demand for cell production such as the epidermis (Cockburn et al., 2022). By contrast, the dependence of late differentiation steps on cell cycle arrest may ensure that highly specialized structures such as the brush border do not form in cells that will divide again.

The mechanisms by which CKI-1 degradation delays EPS-8 and HUM-5 accumulation remain to be investigated. This effect could be due to the delay in cell cycle arrest per se and to the prolonged activity of cyclin-CDK complexes. Notably, cyclin-CDK complexes have been shown to regulate actin-binding proteins. In particular, in *C. elegans* enterocytes, enterohemorrhagic *E. coli* intriguingly induce microvilli effacement and redistribution of ACT-5 to the cytoplasm in a CDK-1 dependent manner. This effect of CDK-1 on ACT-5 involves the phosphorylation of the formin CYK-1 by CDK-1 (Huang et al., 2021). It is therefore plausible that introducing an additional cell cycle modifies actin dynamics and impairs the recruitment of microvilli components through the abnormally late activity of cyclin-CDK complexes. Alternatively, it is possible that the delayed accumulation of EPS-8 and HUM-5 is not directly due to prolonged cell cycle but to the lack of a cell-cycle independent function of CKI-1. It has for instance been shown that the CKI homologue p57^Kip2^ regulates intestinal stem cell fate through the CDK independent regulation of the transcription factor Ascl2 (Creff et al., 2023). Moreover, p27^Kip21^ has also been shown to directly regulate the Rho pathway or actin binding proteins such as cortactin (e.g. (Gui et al., 2014; Jeannot et al., 2017)). Considering our observations that several aspects of intestinal differentiation as well as ELT-2 accumulation are not affected by CKI-1 degradation, we do not favor a model in which CKI-1 would regulate EPS-8 and HUM-5 through transcriptional regulation. We can however not exclude that the loss of CKI-1 induces alterations in the Rho pathway or directly affects some specific actin-binding proteins including EPS-8 and HUM-5.

While our first results indicated a partial effect of cell cycle regulators on intestinal differentiation, we next asked whether differentiation factors reciprocally regulate the ability of intestinal cells to divide. Previous studies have shown that the properties of intestinal cell cycles are tightly linked to the acquisition of intestinal cell fate. Intestinal specification is in particular both necessary and sufficient to control the number of intestinal cell divisions. Pioneering work has indeed shown that isolated E blastomeres divide to produce a 16-20 cell cyst-like structure, indicating that they contain most of the information required to control the number of intestinal cell cycles (Leung et al., 1999). It was also shown that defects in intestinal specification lead to the appearance of supernumerary E descendants, suggesting that specification signals restrict the number of intestinal divisions (Choi et al., 2017; Robertson et al., 2014). It is however noteworthy that several hours separate intestinal specification and cell cycle arrest, raising the question of the signal that may act as a relay between specification factors and exit of the cell cycle. Our observations demonstrate that, although specific differentiation steps, namely polarization and brush border formation, do not trigger cell cycle arrest, the differentiation factors ELT-2 and ELT-7 themselves prevent the occurrence of late intestinal divisions. Our results show that ELT-2 and ELT-7 allow the accumulation of CKI-1 in posterior enterocytes after the E16/E20 transition. In the absence of ELT-2 and ELT-7, the lack of CKI-1 accumulation following the E16 to E20 division of posterior cells is likely to contribute to their ability to reenter the cell cycle, to re-accumulate cyclin B1 and to divide once more.

In mammals several GATA factors have been shown to directly promote the transcription of CKI homologues (Agnihotri et al., 2009; Papetti et al., 2010; Xu et al., 2021), suggesting that ELT-2 and ELT-7 could directly control CKI-1 expression. Nevertheless, although ELT-7 and ELT-2 start to be expressed in all intestinal cells from the E2/E8 stages on (Demouchy et al., 2024; Sommermann et al., 2010), our results strikingly show that they do not inhibit divisions prior to the E20 stage but rather specifically prevent the late occurrence of supernumerary divisions in the most posterior intestinal cells. These observations suggest that additional mechanisms ensure both cell cycle progression during early intestinal development and cell cycle arrest in non-posterior enterocytes. Importantly, we find that the presence of the Hox protein PHP-3 in posterior enterocytes contributes to their ability to undergo supernumerary divisions in the absence of ELT-2 and ELT-7. Very interestingly, the *C. elegans* Hox protein LIN-39 has been shown to be required for cycle progression and to regulate the expression of several *cdk* and *cyclin* genes (Heinze et al., 2023; Roiz et al., 2016). Moreover, constitutive expression of *lin-39* appears to be sufficient to induce overproliferation of sex myoblasts, precocious divisions of Pn.p ventral epidermal cells and prolonged proliferation of vulva cells (Heinze et al., 2023). Hox proteins may thus be critical players in determining the number of cell divisions undergone by each cell type and thus in allowing *C. elegans* to develop according to its typical fixed lineage. Noteworthy, Hox genes are overexpressed in many tumour types where they promote several aspects of tumorigenesis, including cancer cell proliferation (Kopec et al., 2025; Yadav et al., 2024). Altogether, our work and those previous studies point to a broad role for the Hox proteins in cell cycle regulation, which may have important implications in coordinating proliferation and differentiation during both development and tumour formation.

In conclusion, our work reveals the existence of reciprocal interactions between the cell proliferation and cell differentiation programs during the development of the C*. elegans* intestine. We show on one hand that delaying cell cycle arrest hampers the accumulation of late microvilli components. On the other hand, we find that the differentiation factors ELT-2 and ELT-7 are required to prevent late supernumerary divisions in posterior enterocytes. These mechanisms are likely to be important to coordinate intestinal differentiation with intestinal divisions. We however find that not all differentiation steps require prior cell cycle arrest and, reciprocally, that cell cycle arrest in most intestinal cells does not rely on differentiation factors. This study thus underlines that proliferation and differentiation are only partially coupled, suggesting the existence of additional mechanisms ensuring their temporal control.

## Materials and Methods

### Worm strains

Strains were grown on agar plates containing NGM growth media, seeded with *E. coli* (OP50 strain) and maintained at 20°C. The strains used in this study are listed in Table S1.

### New CRISPR strains

CRISPR-CAS9-genome edited CKI-1::AID::BFP strain was generated at the SEGiCel facility (SFR Santé Lyon Est, CNRS UAR 3453, Lyon, France). It was obtained by inserting a GASGASGAS linker, an Auxin Inducible Degron (AID, MPKDPAKPPAKAQVVGWPPVRSYRKNVMVSCQKSSGGPEAAAFVK) (Zhang et al., 2015), a GGGGSGGGG linker and tagBFP2 to the C-terminus of CKI-1.

### Auxin induced degradation

Intestine specific degradation of CKI-1 or PAR-3 was induced by incubating at 20°C embryos expressing CKI-1::AID::BFP (this study) or GFP::AID::PAR-3 (Castiglioni et al., 2020) and the auxin receptor TIR-1 under the control of the *elt-2* promoter (Castiglioni et al., 2020) in 5 mM IAA-AM (acetoxymethyl indole-3-acetic acid, a cell permeable form of auxin which is able to trigger degradation in embryos (Negishi et al., 2019)). The effect of CKI-1 degradation on intestinal cell number (Fig. S2C) and differentiation (Fig.1 and Fig.2) was analysed in early lima bean embryos after 3h auxin treatment, in comma, 1.5-fold and 2-fold embryos after 4h auxin treatment and in 3-fold and 4-fold embryos after 5h auxin treatment. Movies of intestinal divisions (Fig. S2B) were started after 3h auxin treatment. The efficiency of PAR-3 degradation was observed in lima bean to 1.5-fold embryos after 30 min auxin treatment (Fig. S3A), its effect on intestinal nuclei number in 2-fold embryos was observed after 4h auxin treatment.

### RNAi vectors and feeding conditions

RNAi was performed by feeding worms with RNAi clones on 1 mM IPTG plates during 24 h (*act-5* RNAi) or 48 h (*elt-2* RNAi). L4440 was used as a control (Timmons and Fire, 1998). *act-5* RNAi was performed with the appropriate clone (sjj_T25C8.2, III-6D08) from the Ahringer-Source Bioscience library (Kamath et al., 2003). For *elt-2* RNAi, a 555bp fragment amplified from the *elt-2* genomic sequence (oligos F-tacaacaaatcaatacagg and R-ataaaagttgatcacgtacc) was cloned into a Gateway compatible L4440 vector.

### Microscopy and time-lapse recordings

For still imaging of live embryos, embryos were mounted on 2% agarose pads in a drop of M9 medium enriched in OP50 bacteria to immobilize embryos. Standard confocal images were acquired with a SP8 confocal microscope (Leica) equipped with a HC Plan-Apo 63×, 1.4 NA objective and the LAS AF software. Z-stacks were acquired with 300 to 500 nm steps. Images of autofluorescent gut granules (Fig.1B) were obtained by illuminating embryos with a 405 nm diode. Images used to quantify the nuclear levels of ELT-2::mNG (Fig.1A) and the apical accumulation of microvilli components (Fig.2) were acquired in the photocounting mode and with a number a line accumulations chosen to be sufficient for detecting the signal and avoiding saturation. For EPS-8::mNG and HUM-5::mNG, the strong signal increase occurring during development forced us to use different numbers of accumulations for different embryonic stages. The number of accumulations used were then taken into account for quantification of signal intensity (see below) and are indicated on the corresponding figures (Fig.2C and 2D). Superresolution images were acquired with a LSM 880 microscope (Zeiss), equipped with a Plan-Apo 63×, 1.4 NA objective and the Zen Black software. The Airy Scan module was used to obtain superresolution images.

For time-lapse experiments, adult hermaphrodites were dissected in M9 medium. Embryos were then transferred to a 4 µL drop of M9 medium on a 35 mm glass bottom dish (poly-D-lysine coated, 14 mm microwell, No 1.5 coverglass, MatTek). Excess of M9 was then removed to ensure adhesion of embryos to poly-D-Lysine and 2 mL of mineral oil (Light Oil (neat), Sigma) was used to cover the drop of M9. Recordings were performed on a Leica DMi8 spinning disc microscope equipped with a HCX Plan-Apo 63×, 1.4 NA objective and a photometrics Evolve EMCCD camera. The setup was controlled by the Inscoper Imaging Suite. Embryos were maintained at 23°C during recordings. Images were acquired at 3 minutes (Fig.4, 5, 6, S5, S6, S7, S8) or 4 minutes (Fig. S2B) intervals. Z-stacks were acquired with 400 nm steps.

### Image analysis and quantifications

Images were assembled for illustration using the Fiji and Inkscape 1.1 softwares. All embryos are oriented with the anterior pole to the left. Max intensity z-projections were obtained with Fiji. Quantifications of ELT-2 nuclear levels were performed with Fiji by measuring and averaging fluorescence intensity in all intestinal nuclei. Quantifications of ERM-1, FLN-2, EPS-8 and HUM-5 apical levels were performed with Fiji, by measuring mean fluorescence intensity along a segmented line covering the whole intestine and then dividing the measured fluorescence by the number of line accumulations used for each stage. Images from movies in Fig.4, 5, 6, S5, S6, S7 and S8 have been post-treated with a 3D Gaussian filter (X sigma=2, Y sigma=2, Z sigma=2, Fiji).

### Statistical analysis

Statistical analysis and graphic representations were performed with the Graphpad Prism 10.0.3 software. Graphs show all individual values (dots), mean and standard deviation (black lines). Details of the statistical tests used are indicated in each figure legend. Two-tailed unpaired Student’s t tests were performed when data passed the Shapiro and Kolmogorov-Smirnov normality tests and when sample variance was equal; two-tailed Welch’s t tests were performed when data passed the Shapiro and Kolmogorov-Smirnov normality tests and when sample variance was unequal; two-tailed non-parametric Mann-Whitney tests were performed when data did not pass the Shapiro or Kolmogorov-Smirnov normality test.

## Acknowledgments

We thank M. Boxem, J. Sepers and H. Pires for kindly sharing their GFP::degron::PAR-3 strain prior to publication as well as M. Boxem, V. Gobel, K. Kemphues, M. Maduro, M. Soto and the *Caenorhabditis* Genetics Center (funded by the National Institute of Health Office of Research Infrastructure Programs, P40 OD010440) for providing worm strains. The CKI-1::degron strain was generated by the SEGiCel Facility (SFR Santé Lyon Est, CNRS UAR 3453, Lyon, France). M. Kanemaki provided us with modified auxin. We thank the Michaux and Pecreaux labs as well as L. Pintard, F. Pituello and L. Bataillé for valuable discussions. Imaging was performed at the Microscopy Rennes imaging Center (MRiC Photonics, Biosit, Rennes, France), a member of the national infrastructure France-BioImaging supported by the French National Research Agency (ANR-10-INBS-04).

## Author contributions

Conceptualization: J.D., G.M., A.P.; Methodology: J.D., G.M., A.P.; Validation: J.D., H.S., A.P.; Formal analysis: J.D., H.S., A.P.; Investigation: J.D., H.S., A.P.; Writing – original draft preparation: J.D., A.P.; Writing – Review and editing: J.D., H.S., G.M., A.P.; Vizualization: J.D., A.P.; Supervision: G.M., A.P.; Project administration: G.M.; Funding Acquisition: G.M., A.P.

---

**Fig. S1:**
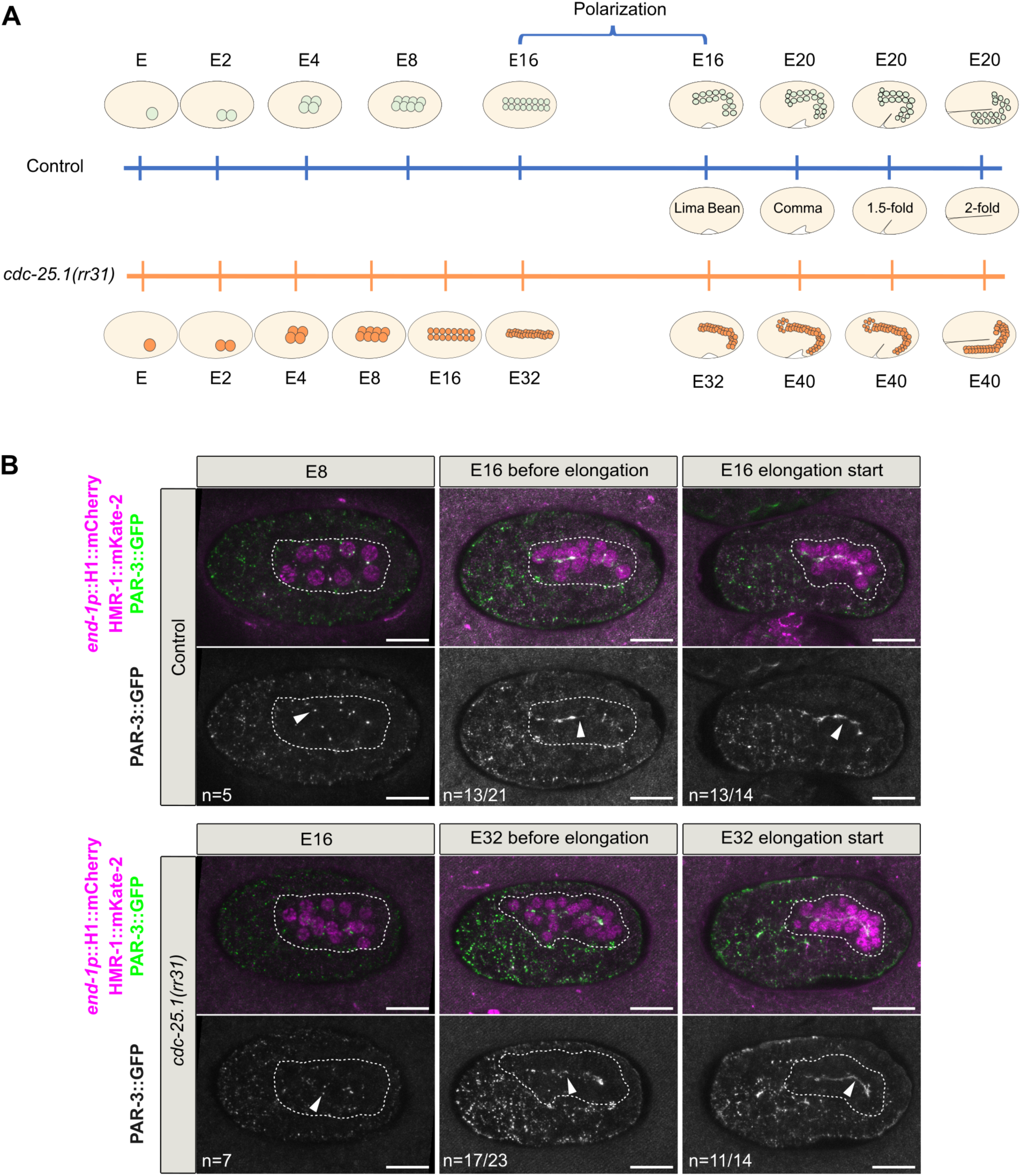
Acceleration of intestinal cell cycles triggered by a *cdc-25.1* gain of function mutation does not lead to precocious polarization. **A.** Schematic representation of intestinal divisions in control and *cdc-25.1(rr31)* embryos. The E16 stage is reached before the start of elongation while 2-fold embryos are at the E20 stage. Polarization occurs at the E16 stage. In *cdc-25.1(rr31)* embryos, enterocytes divide faster and one more time than in control embryos, such that the intestine reaches the E16 stage approximately at the same time as the E8 stage in control embryos and the E32 stage before elongation initiation (Demouchy et al., 2024; Kostić and Roy, 2002). **B.** Maximum Z-projections from confocal images of control and *cdc-25.1(rr31)* embryos expressing PAR-3::GFP (green), HMR-1::mKate2 (magenta) and the intestinal nuclear marker *end1p::*H1::mCherry (magenta). In control (n=5) and *cdc-25.1(rr31)* (n=7) embryos, non-polarized PAR-3 puncta (arrowheads) are visible at the E8 and E16 stage, respectively. At the E16 (control) and E32 (*cdc-25.1(rr31))* stages prior to elongation, most embryos display discontinuous apical foci (arrowheads, n=13/21 control and 17/23 *cdc-25.1(rr31),* images shown). Other embryos display a continuous apical PAR-3 line (n=8/21 control and 6/23 *cdc-25.1(rr31),* not shown). At the beginning of elongation PAR-3 forms a continuous apical line (arrowheads) in most control (E16, n=13/14) and *cdc-25.1(rr31)* (E32, n=11/14) embryos. Other embryos display a discontinuous apical line (n=1/14 control and 3/14 *cdc-25.1(rr31),* not shown). White dashed lines highlight embryo shape. Scale bars: 10 µm.

**Fig. S2:**
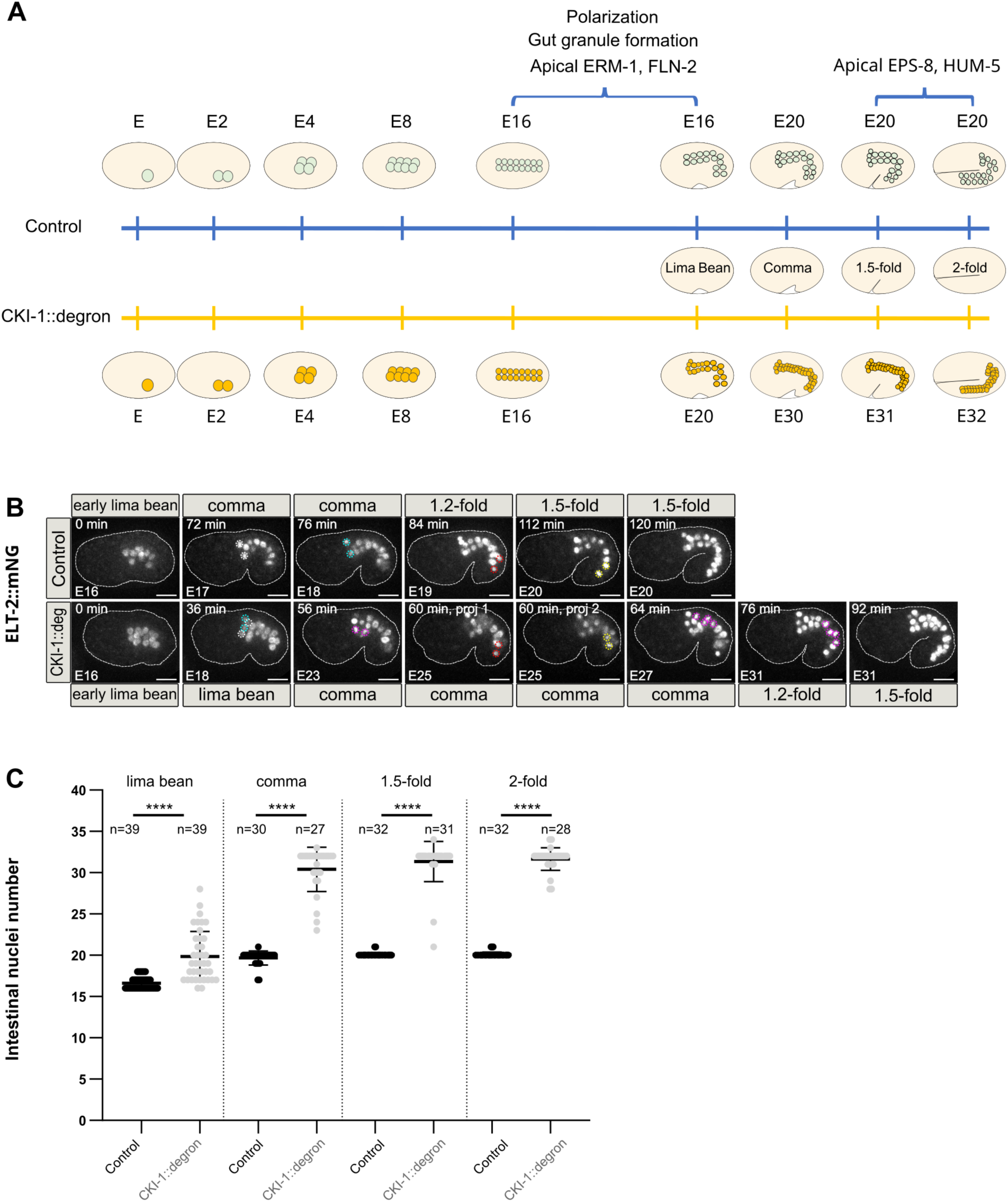
Loss of CKI-1 leads to the appearance of a late supplementary enterocyte division. **A.** Schematic representation of intestinal divisions in control embryos and in embryos lacking CKI-1. Until the E16 stage, enterocytes divide at the same speed in both types of embryos; in embryos lacking CKI-1, all intestinal cells then undergo an additional late division after the E16 stage so that embryos finally display 32 cells (Kostić and Roy, 2002). In control embryos, polarization, gut granule formation and apical accumulation of the first microvilli components (such as ERM-1 and FLN-2) occur at the E16 stage while apical accumulation of later microvilli components (EPS-8, HUM-5) occurs between the 1.5 and 2-fold stage. **B.** Maximum Z-projections from confocal images of auxin-treated embryos expressing the intestinal nuclear marker ELT-2::mNG and either *elt-2p::*TIR-1 (Control, n=8) or both *elt-2p::*TIR-1 and CKI-1::degron::BFP (CKI-1::deg, n=9) and recorded between the beginning of elongation and the 1.5-fold stage. Dotted circles highlight the daughter cells obtained just after the division of the two cells from the first (white/cyan) or last (red/yellow) intestinal ring as well as some of the cells resulting from the abnormal divisions of central enterocytes in CKI-1::degron embryos (magenta). White dashed lines highlight embryo shape. Scale bars: 10 µm. **C.** Quantification of intestinal nuclei number in auxin-treated embryos expressing the intestinal nuclear marker ELT-2::mNG and either *elt-2p::*TIR-1 (Control) or both *elt-2p::*TIR-1 and CKI-1::degron::BFP (CKI-1::deg) and observed at the indicated embryonic stages. \*\*\*\**P*<0.0001, Mann-Whitney test.

**Fig. S3:**
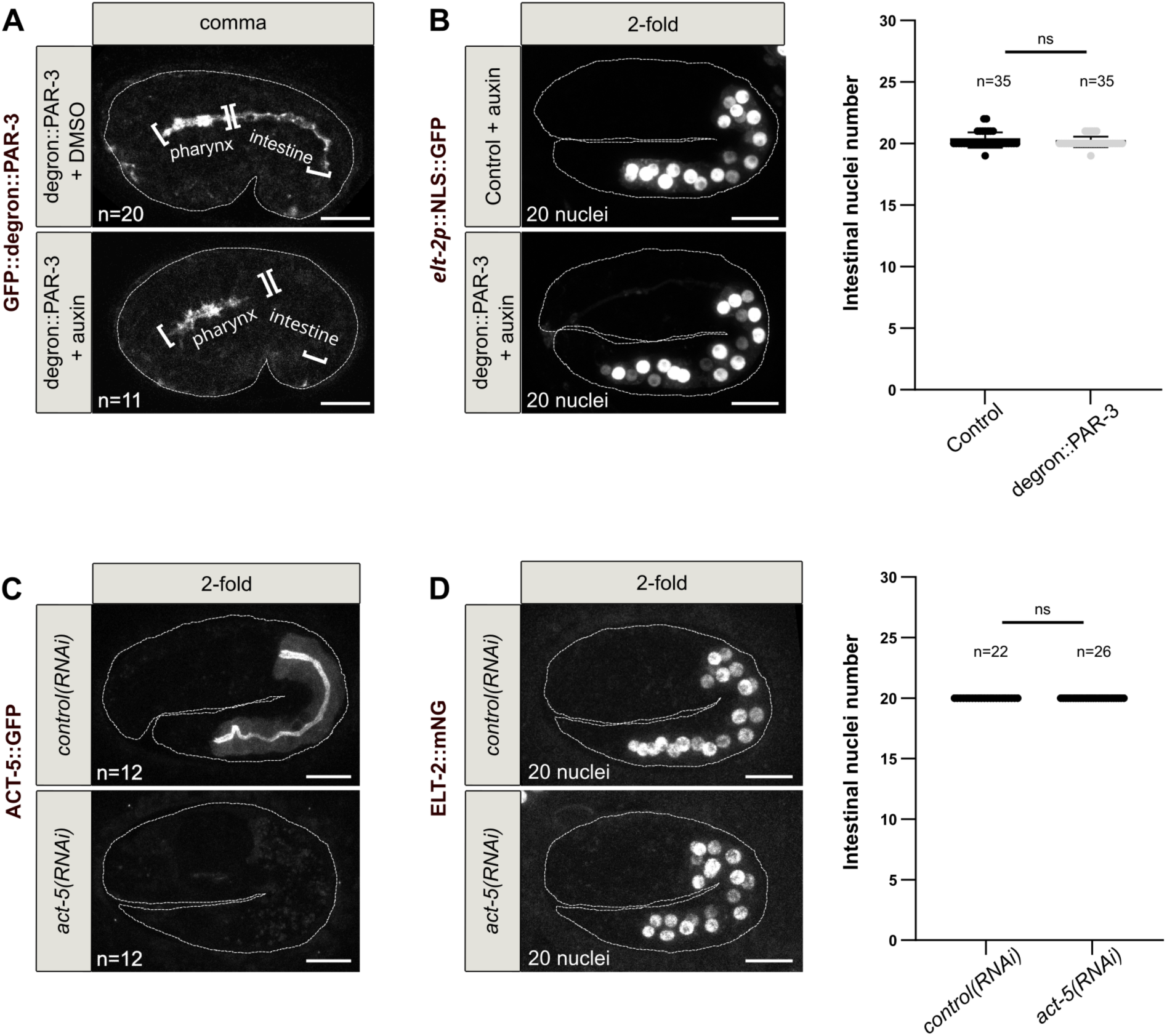
Intestinal polarization and brush border formation are not required to arrest enterocyte divisions. **A.** Maximum Z-projections from confocal images of comma embryos expressing GFP::degron::PAR-3 and *elt-2p::*TIR-1 and treated 30 min with DMSO or auxin. In the presence of auxin, PAR-3 is specifically degraded in intestinal cells. **B.** Maximum Z-projections from confocal images of 2-fold embryos expressing the intestinal nuclear marker *elt-2p::*NLS::GFP and either *elt-2p::*TIR-1 (Control) or both *elt-2p::*TIR-1 and GFP::degron::PAR-3 after 4h auxin treatment and quantification of the number of intestinal nuclei in those embryos. ns: non-significant (*P*>0.05), Mann-Whitney test. **C.** Maximum Z-projections from confocal images of 2-fold embryos expressing ACT-5::GFP and treated with control and *act-5* RNAi. Upon *act-5* RNAi, all embryos showed an absence of ACT-5::GFP signal. **D.** Maximum Z-projections from confocal images of 2-fold embryos expressing the intestinal nuclear marker ELT-2::mNG and treated with control or *act-5* RNAi and quantification of the number of intestinal nuclei in those embryos. ns: non-significant (*P*>0.05), Mann-Whitney test. In A to D, white dashed lines highlight embryo shape. Scale bars: 10 µm.

**Fig. S4:**
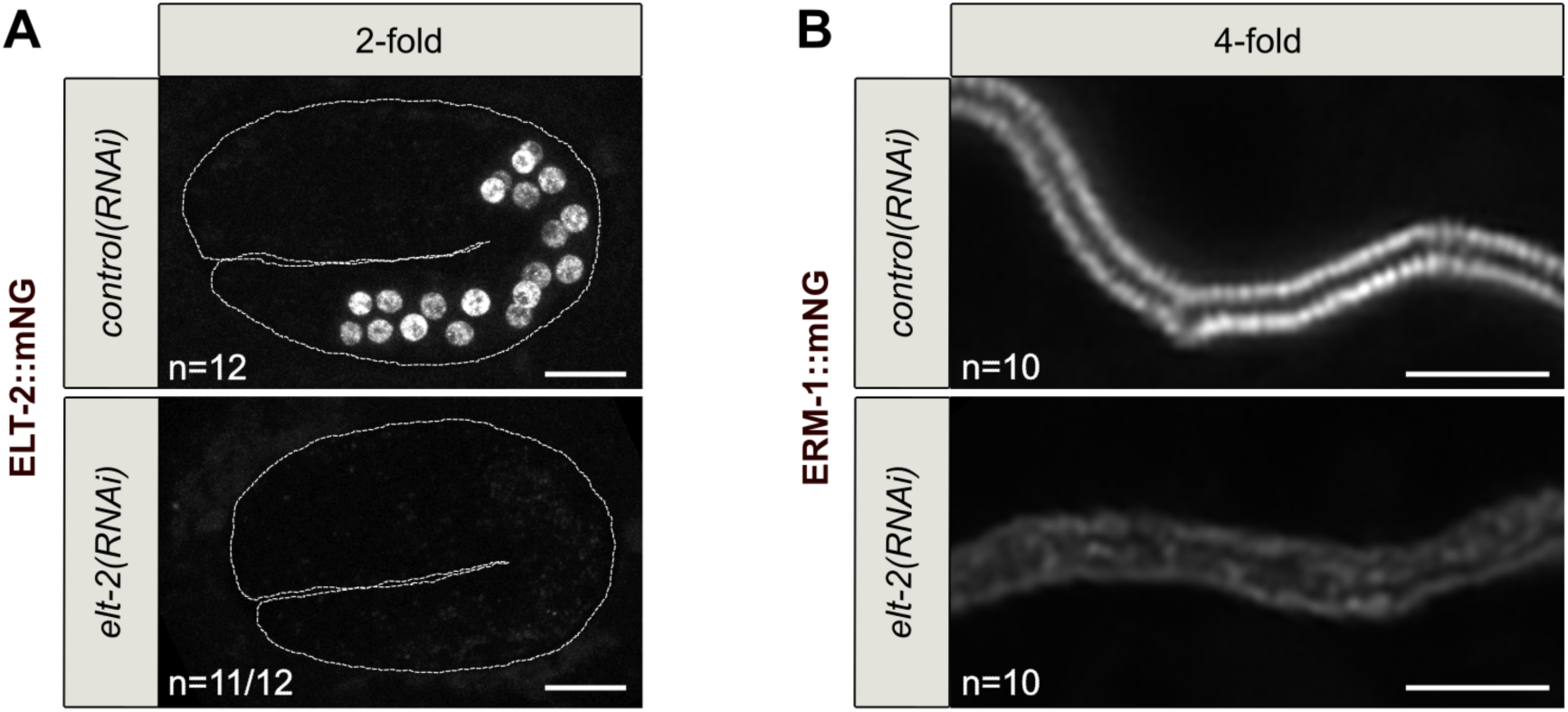
Loss of ELT-2 leads to strong intestinal defects. **A.** Maximum Z-projections from confocal images of 2-fold embryos expressing the intestinal nuclear marker ELT-2::mNG and treated with control or *elt-2* RNAi. Upon *elt-2* RNAi, 11/12 embryos showed a very strong reduction of ELT-2::mNG signal; some residual ELT-2::mNG signal was observed in some enterocytes in 1/12 embryo. White dashed lines highlight embryo shape. Scale bars: 10 µm. **B.** Super-resolution confocal images of the intestine in 4-fold embryos expressing ERM-1::mNG and treated with control or *elt-2* RNAi. In control embryos, the intestine is characterized by a well-defined brush border while *elt-2* RNAi treated embryos display a poorly structured brush border. Scale bars: 2 µm.

**Fig. S5:**
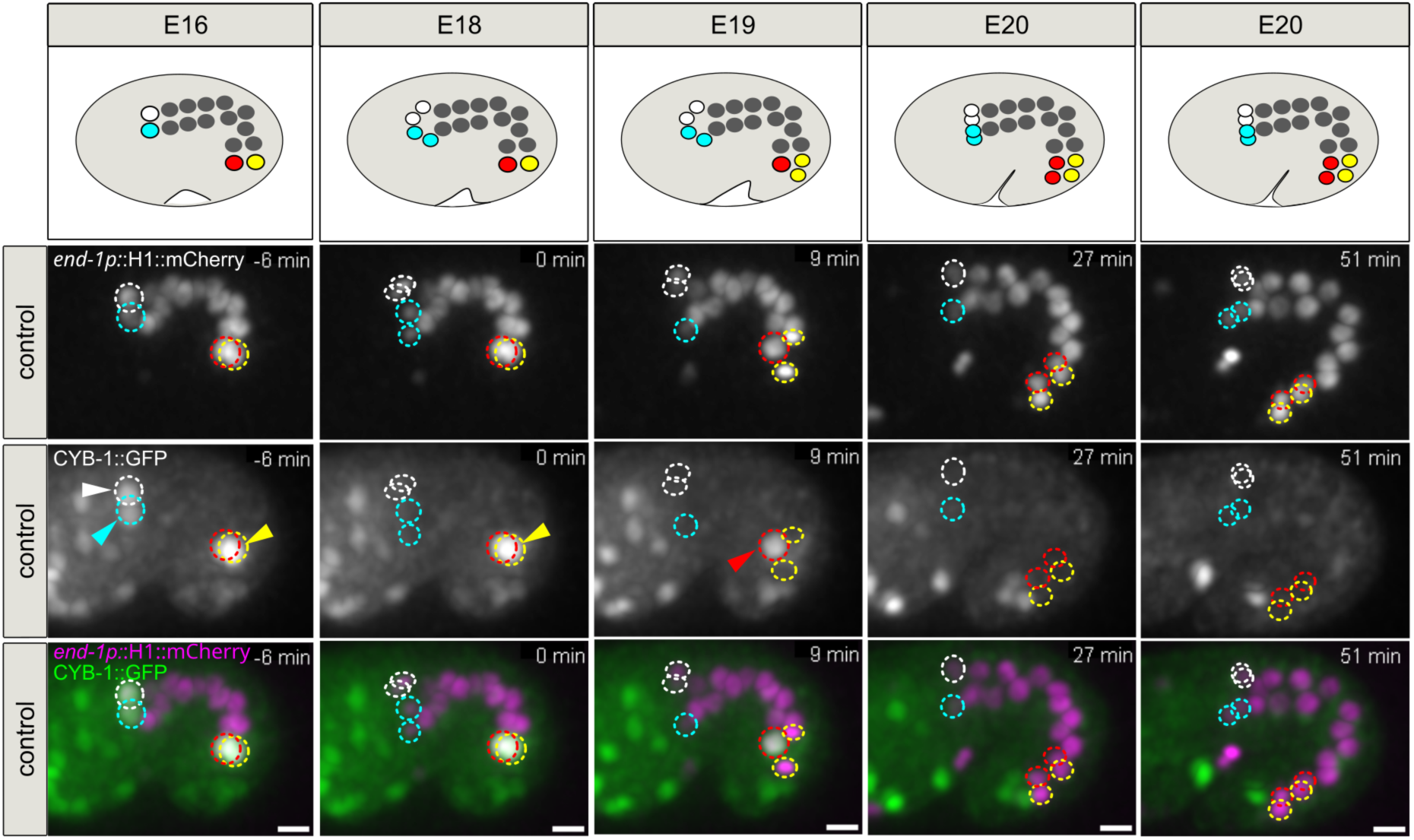
Expression pattern of CYB-1::GFP in control embryos. Schematic representations and maximum Z-projections from confocal images showing the enterocytes of a control embryo expressing the intestinal nuclear marker *end1p::*H1::mCherry and CYB-1::GFP and recorded between the E16 and E20 stage. White/cyan and red/yellow circles highlight the enterocytes from the first and last intestinal ring, respectively, before (E16) and after (E17 to E20) their division. The first division of the E16 to E20 transition was used as reference for division timing (t=0). In this embryo, the two first ring divisions occur at t=0 (white and cyan nuclei). Last ring divisions occur at t=9 min (yellow nuclei) and t= 27 min (red nuclei). CYB-1::GFP accumulates specifically in the first and last ring enterocytes prior to their division (arrowheads) and does not reappear in their daughter cells. Scale bars: 5 µm.

**Fig. S6:**
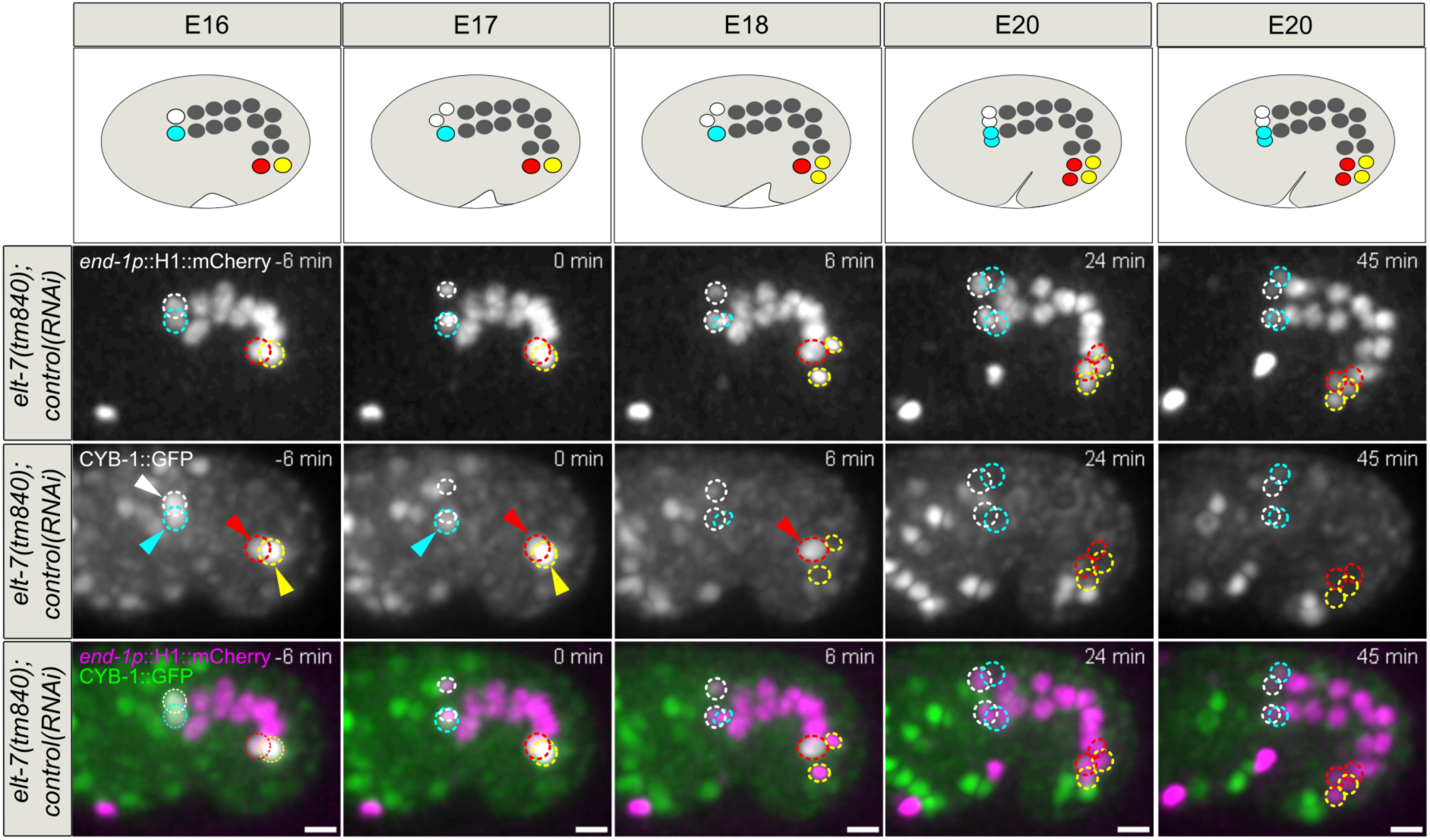
Expression pattern of CYB-1::GFP in *elt-7(tm840)* embryos. Schematic representations and maximum Z-projections from confocal images showing the enterocytes of a *elt-7(tm840)* embryo treated with control RNAi, expressing the intestinal nuclear marker *end1p::*H1::mCherry and CYB-1::GFP and recorded between the E16 and E20 stage. White/cyan and red/yellow circles highlight the enterocytes from the first and last intestinal ring, respectively, before (E16) and after (E17 to E20) their division. The first division of the E16 to E20 transition was used as reference for division timing (t=0). In this embryo, first ring divisions occur at t=0 (white nuclei) and between t=6 to t= 24 min (cyan nuclei). Last ring divisions occur at t=6 min (yellow nuclei) and between t=6 to t= 24 min (red nuclei). CYB-1::GFP accumulates specifically in the first and last ring enterocytes prior to their division (arrowheads) and does not reappear in their daughter cells. Scale bars: 5 µm

**Fig. S7:**
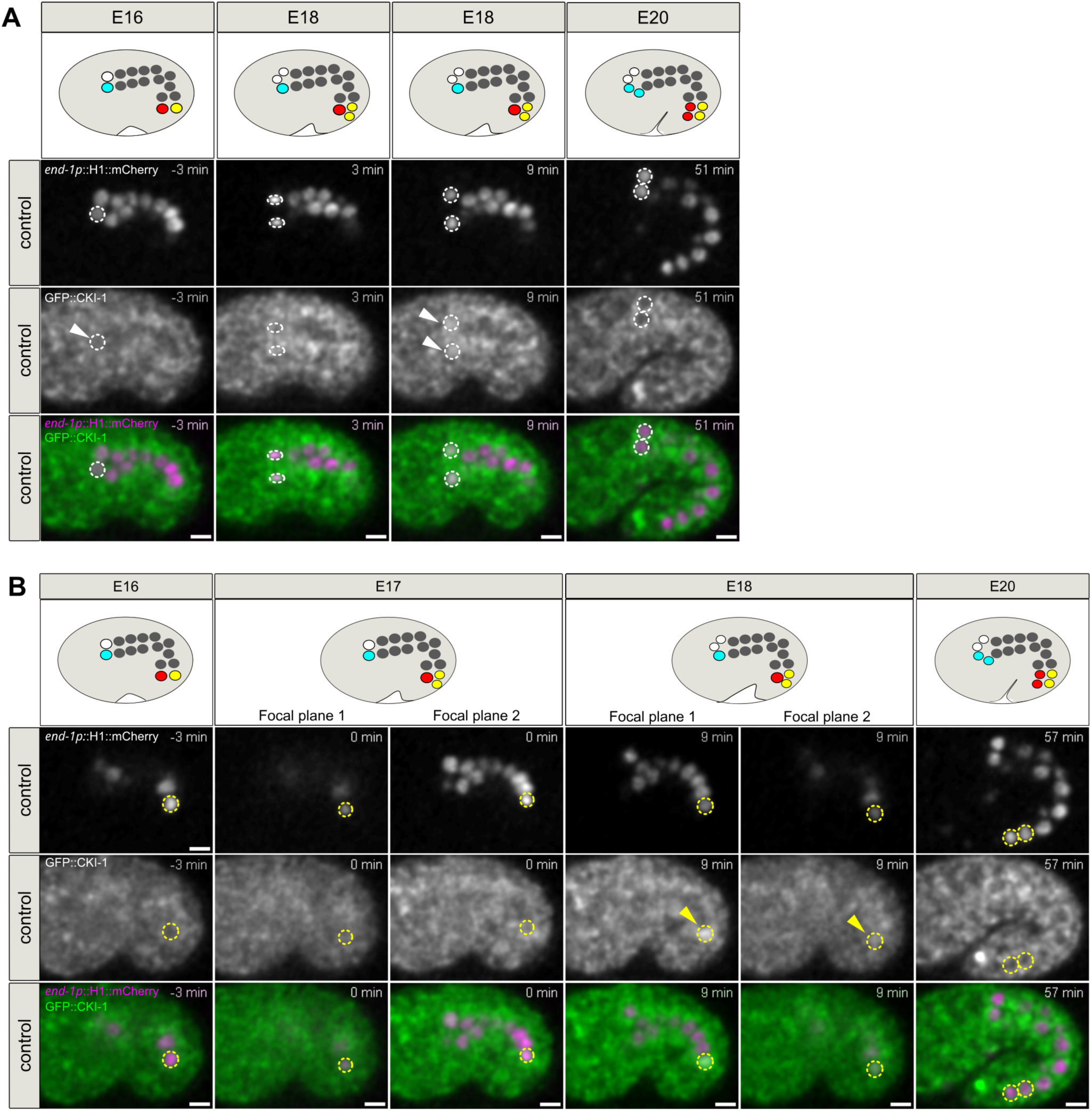
Expression pattern of GFP::CKI-1 in control embryos. **A-B.** Schematic representations and confocal images showing the enterocytes of a control embryo expressing the intestinal nuclear marker *end1p::*H1::mCherry and GFP::CKI-1 and recorded between the E16 and E20 stage. The same embryo is shown in A and B; focal planes were chosen to display the anterior (A) or posterior (B) cell divisions. White/cyan and red/yellow circles highlight the nuclei of one cell of the first and last intestinal ring enterocytes, respectively, before (E16) and after (E17 to E20) their division. The first division of the E16 to E20 transition was used as reference for division timing (t=0). In this embryo, one first ring division (A) occurs at t=3 min (white nucleus). Division of the other anterior cell (cyan nucleus) occurs in a different focal plane and is not shown here. GFP::CKI-1 starts to be slightly visible in the white mother cell just prior to division (t=-3 min, arrowhead) and accumulates in the nucleus of the two white daughter cells after the division (t=9 min, arrowheads) and then progressively disappears until it is not any longer visible (t=51 min). One last ring division (B) occurs at t=0 (yellow nuclei, 2 focal planes are shown to make the two daughter nuclei visible). Division of the other posterior cell (red nucleus) occurs in a different focal plane and is not shown here. GFP::CKI-1 accumulates in the nucleus of the two yellow daughter cells just after the division (t=9 min, arrowheads, 2 focal planes are shown to make the yellow daughter nuclei visible) and then progressively disappears until it is not any longer visible (t=57 min). Scale bars: 5 µm.

**Fig. S8:**
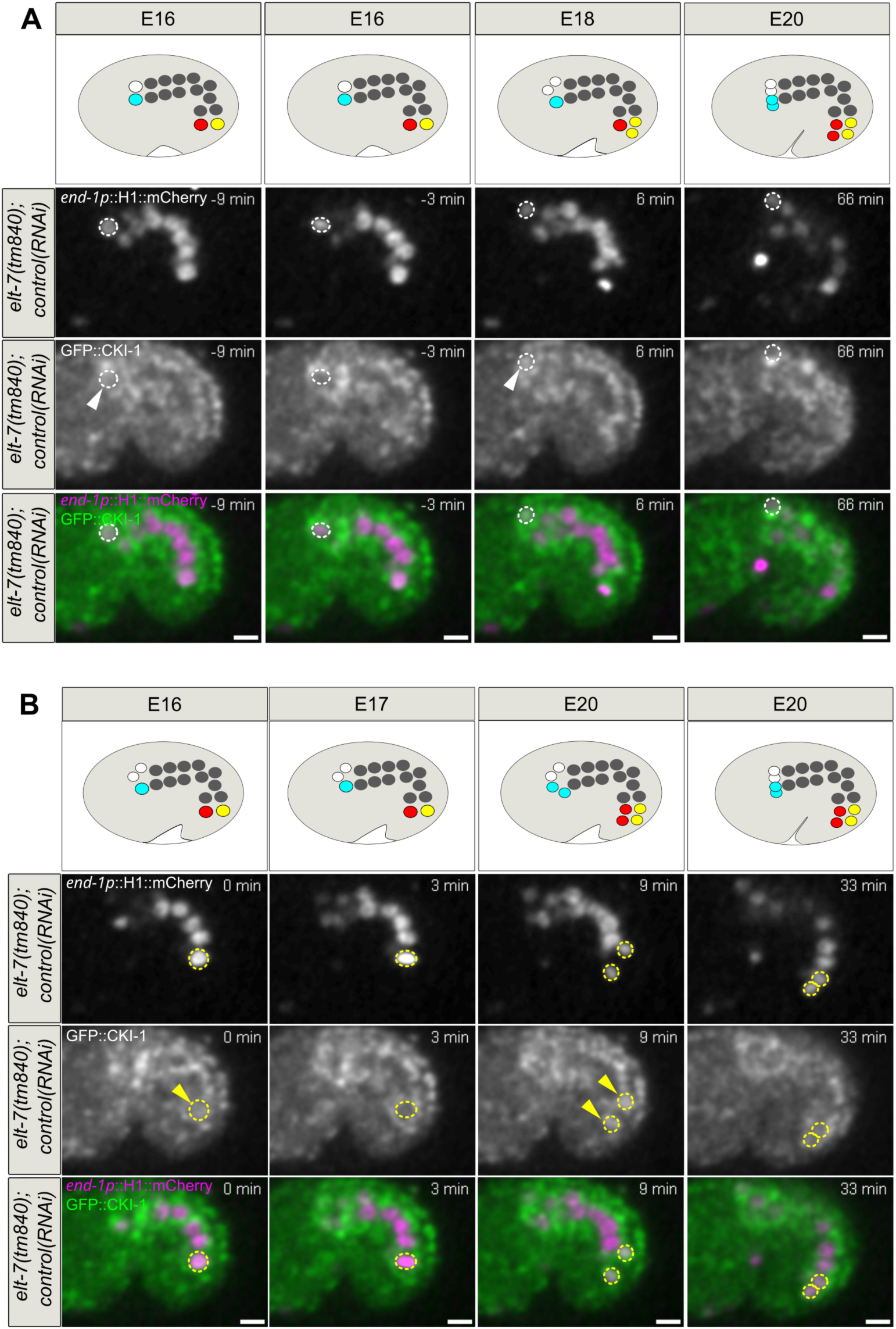
Expression pattern of GFP::CKI-1 in *elt-7(tm840)* embryos. Schematic representations and confocal images showing the first and last rings of a *elt-7(tm840)* embryo treated with control RNAi, expressing the intestinal nuclear marker *end1p::*H1::mCherry and GFP::CKI-1 and recorded between the E16 and E20 stage. The same embryo is shown in A and B; focal planes were chosen to display the anterior (A) or posterior (B) cell divisions. White/cyan and red/yellow circles highlight the nuclei of one cell of the first and last intestinal ring enterocytes, respectively, before (E16) and after (E17 to E20) their division. The first division of the E16 to E20 transition was used as reference for division timing (t=0). In this embryo, one of the first ring divisions (A) occurs at t=0 (white nuclei, only one daughter cell visible in the chosen focal plane). Division of the other anterior cell (cyan nucleus) occurs in a different focal plane and is not shown here. GFP::CKI-1 starts to be slightly visible in the white mother cell nucleus prior to division (t=-9 min, arrowhead) and accumulates in the nucleus of the two white daughter cells after the division (t=6 min, arrowhead, only one daughter cell visible) and then progressively disappears until it is not any longer visible (t=66 min). One of the last ring divisions (B) occurs between t=3 and t=6 min (yellow nuclei). Division of the other posterior cell (red nucleus) occurs in a different focal plane and is not shown here. GFP::CKI-1 starts to be slightly visible in the yellow mother nucleus prior to division (t=0 min, arrowhead) and accumulates in the nucleus of the two yellow daughter cells after the division (t=9 min, arrowheads) and then progressively disappears until it is not any longer visible (t=33 min). Scale bars: 5 µm.

**Table S1:** *C. elegans* strains used in this study.

| Strain | Complete genotype | Source | Figures |
| --- | --- | --- | --- |
| FL781 | *mibls48[elt-2p::TIR-1::tagBFP2-Lox511::tbb-2-3'UTR, IV:5014740-5014802(cxTi10882 site)] IV; elt-2(bab354[elt-2::mNG]) X* | This study<br>*mibls48* from BOX273 (Boxem lab (Castiglioni et al., 2020))<br>*elt-2(bab354)* from FL660 (Demouchy et al., 2024) | Fig.1A-B<br>Fig.S2B-C |
| FL780 | *cki-1(bab393[cki-1::AID::tagBFP2]) II; mibls48[elt-2p::TIR-1::tagBFP2-Lox511::tbb-2-3'UTR, IV:5014740-5014802(cxTi10882 site)] IV; elt-2(bab354[elt-2::mNG]) X* | This study<br>*cki-1(bab393)* from FL733 (this study) | Fig.1A-B<br>Fig.S2B-C |
| FL754 | *par-6(it310[par-6::GFP]) I; stls10220[end-1p::his-24::mCherry::let-858 3' UTR + unc-119(+)] III; mibls48[elt-2p::TIR-1::tagBFP2-Lox511::tbb-2-3'UTR, IV:5014740-5014802(cxTi10882 site)] IV* | This study<br>*par-6(it310)* from KK1248 (Kemphues lab)<br>*stls10220* from RW10234 (Waterston lab, via CGC) | Fig.1C |
| FL778 | *par-6(it310[par-6::GFP]) I; cki-1(bab393[cki-1::AID::tagBFP2]) II; stls10220[end-1p::his-24::mCherry::let-858 3' UTR + unc-119(+)] III; mibls48[elt-2p::TIR-1::tagBFP2-Lox511::tbb-2-3'UTR, IV:5014740-5014802(cxTi10882 site)] IV* | This study | Fig.1C |
| FL748 | *erm-1(bab59[erm-1::mNG^3xFlag]) I; stls10220[end-1p::his-24::mCherry::let-858 3' UTR + unc-119(+)] III; mibls48[elt-2p::TIR-1::tagBFP2-Lox511::tbb-2-3'UTR, IV:5014740-5014802(cxTi10882 site)] IV* | This study<br>*erm-1(bab59)* from FL378 (Bidaud-Meynard et al., 2019) | Fig.2A |
| FL763 | *erm-1(bab59[erm-1::mNG^3xFlag]) I; cki-1(bab393[cki-1::AID::tagBFP2]) II; stls10220[end-1p::his-24::mCherry::let-858 3' UTR + unc-119(+)] III; mibls48[elt-2p::TIR-1::tagBFP2-Lox511::tbb-2-3'UTR, IV:5014740-5014802(cxTi10882 site)] IV* | This study | Fig.2A |
| FL574 | *mibls48[elt-2p::TIR-1::tagBFP2-Lox511::tbb-2-3'UTR, IV:5014740-5014802(cxTi10882 site)] IV; fln-2(jme-19)[fln-2::mVenus]) X* | This study<br>*fln-2(jme-19)* from FBR222 (Robin lab, (Bidaud-Meynard et al., 2021)) | Fig.2B |
| FL782 | *cki-1(bab393[cki-1::AID::tagBFP2]) II; mibls48[elt-2p::TIR-1::tagBFP2-Lox511::tbb-2-3'UTR, IV:5014740-5014802(cxTi10882 site)] IV; fln-2(jme-19)[fln-2::mVenus]) X* | This study | Fig.2B |
| FL755 | *stls10220[end-1p::his-24::mCherry::let-858 3' UTR + unc-119(+)] III; eps-8(bab140[eps-8::mNG]) IV mibls48[elt-2p::TIR-1::tagBFP2-Lox511::tbb-2-3'UTR, IV:5014740-5014802(cxTi10882 site)] IV* | This study<br>*eps-8(bab140)* from FL290 (Bidaud-Meynard et al., 2021) | Fig.2C |
| FL764 | *cki-1(bab393[cki-1::AID::tagBFP2]) II; stls10220[end-1p::his-24::mCherry::let-858 3' UTR + unc-119(+)] III; eps-8(bab140[eps-8::mNG]) IV mibls48[elt-2p::TIR-1::tagBFP2-Lox511::tbb-2-3'UTR, IV:5014740-5014802(cxTi10882 site)] IV* | This study | Fig.2C |
| FL597 | *hum-5(bab189[hum-5::mNG]) III; mibls48[elt-2p::TIR-1::tagBFP2-Lox511::tbb-2-3'UTR, IV:5014740-5014802(cxTi10882 site)] IV* | This study<br>*hum-5(bab189)* from FL579 (outcrossed from MCP189 (Bidaud-Meynard et al., 2021)) | Fig.2D |
| FL773 | *cki-1(bab393[cki-1::AID::tagBFP2]) II; hum-5(bab189[hum-5::mNG])III; mibls48[elt-2p::TIR-1::tagBFP2-Lox511::tbb-2-3'UTR, IV:5014740-5014802(cxTi10882 site)] IV* | This study | Fig.2D |
| FL630 | *stls10220[end-1p::his-24::mCherry::let-858 3' UTR + unc-119(+)] III* | This study | Fig.3A-B<br>Fig.7B |
| FL734 | *stls10220[end-1p::his-24::mCherry::let-858 3' UTR + unc-119(+)] III; elt-7(tm840) V* | This study<br>*elt-7(tm840)* from MS433 (Mitani lab, via Maduro lab, (McGhee et al., 2007)) | Fig.3A-B<br>Fig.7B |
| FL747 | *stls10220[end-1p::his-24::mCherry::let-858 3' UTR + unc-119(+)] III; elt-7(tm840) V zuls70[end-1p::GFP::CAAX] V* | This study<br>*zuls70* from MS2293 (Maduro lab (Choi et al., 2017), originally from Nance lab (Wehman et al., 2011)) | Fig.3C<br>Fig.4 |
| FL760 | *stls10220[end-1p::his-24::mCherry::let-858 3' UTR + unc-119(+)] III; cyb-1(tn1806[cyb-1::GFP::tev::3xflag]) IV; elt-7(tm840) V* | This study<br>*cyb-1(tn1806)* from DG4569 (Greenstein lab, via CGC (Spike et al., 2022)) | Fig.5<br>Fig.S6 |
| FL758 | *cki-1(bmd132[GFP::LoxP::cki-1::3xFLAG]) II; stls10220[end-1p::his-24::mCherry::let-858 3' UTR + unc-119(+)] III; elt-7(tm840) V* | This study<br>*cki-1(bmd132)* from DQM455 (Matus lab, via CGC (Adikes et al., 2020)) | Fig.6<br>Fig.S8 |
| FL777 | *nob-1(syb2679[nob-1::GFP]) III stls10220[end-1p::his-24::mCherry::let-858 3' UTR + unc-119(+)] III* | This study<br>*nob-1(syb2679)* from PHX2679 (Zacharias lab, via CGC, (Murray et al., 2022)) | Fig.7A |
| FL775 | *php-3(st12213[php-3::TY1::EGFP::3xFLAG]) III stls10220[end-1p::his-24::mCherry::let-858 3' UTR + unc-119(+)] III* | This study<br>*php-3(st12213)* from RW12213 (Waterston lab, via CGC) | Fig.7A |
| FL794 | *php-3(ok919) III stls10220[end-1p::his-24::mCherry::let-858 3' UTR + unc-119(+)] III* | This study<br>*php-3(ok919)* from RB998 (*C. elegans* Gene knockout consortium, via CGC, (The *C. elegans* Deletion Mutant Consortium, 2012)) | Fig.7B |
| FL795 | *php-3(ok919) III stls10220[end-1p::his-24::mCherry::let-858 3' UTR + unc-119(+)] III; elt-7(tm840) V* | This study | Fig.7B |
| FL677 | *hmr-1(cp57[hmr-1::mKate2]) I; par-3(it298[par-3::gfp]) III stls10220[end-1p::his-24::mCherry::let-858 3' UTR + unc-119(+)] III* | This study<br>*hmr-1(cp57)* from OX704 (Soto lab (Sasidharan et al., 2018); originally from LP238, Goldstein lab)<br>*par-3(it298)* from KK1216 (Kempthues lab) | Fig.S1B |
| FL676 | *cdc-25.1(rr31) hmr-1(cp57[hmr-1::mKate2]) I; par-3(it298[par-3::gfp]) III stls10220[end-1p::his-24::mCherry::let-858 3' UTR + unc-119(+)] III* | This study<br>*cdc-25.1(rr31)* from MR142 (Roy lab, via CGC (Kostić and Roy, 2002)) | Fig.S1B |
| FL511 | *par-3(mib68[eGFP-Lox2272::aid::par-3b+eGFP(nolntrons)-LoxP::aid::par-3g]) III; mibls48[elt-2p::TIR-1::tagBFP2-Lox511::tbb-2-3'UTR, IV:5014740-5014802(cxTi10882 site)] IV* | This study<br>*par-3(mib68)* from BOX290 (Boxem lab (Castiglioni et al., 2020)) | Fig.S3A |
| FL607 | *hmr-1(cp57[hmr-1::mKate2]) I; mibls48[elt-2p::TIR-1::tagBFP2-Lox511::tbb-2-3'UTR, IV:5014740-5014802(cxTi10882 site)] IV; rrls1[elt-2p::NLS::GFP::LacZ::unc-54 3'UTR + unc-119(+)] X* | This study<br>*rrls1* from MR142 (Roy lab, via CGC (Kostić and Roy, 2002)) | Fig.S3B |
| FL605 | *hmr-1(cp57[hmr-1::mKate2]) I; par-3(mib68[eGFP-Lox2272::aid::par-3b+eGFP(nolntrons)-LoxP::aid::par-3g]) III; mibls48[elt-2p::TIR-1::tagBFP2-Lox511::tbb-2-3'UTR, IV:5014740-5014802(cxTi10882 site)] IV; rrls1[elt-2p::NLS::GFP::LacZ::unc-54 3'UTR + unc-119(+)] X* | This study | Fig.S3B |
| VJ268 | *fgEx12[act-5p::act-5::GFP + unc-119(+)] II* | Gobel lab (Zhang et al., 2012) | Fig.S3C |
| FL660 | *elt-2(bab354[elt-2::mNG]) X* | (Demouchy et al., 2024) | Fig.S3D<br>Fig.S4A |
| FL378 | *erm-1(bab59[erm-1::mNG^3xFlag]) I* | (Bidaud-Meynard et al., 2019) | Fig.S4B |
| FL752 | *stls10220[end-1p::his-24::mCherry::let-858 3' UTR + unc-119(+)] III; cyb-1(tn1806[cyb-1::GFP::tev::3xflag]) IV* | This study | Fig.S5 |
| FL709 | *stls10220[end-1p::his-24::mCherry::let-858 3' UTR + unc-119(+)] III; cki-1(bmd132[GFP::LoxP::cki-1::3xFLAG]) II* | This study | Fig.S7 |

